# Genomic Insights into the Demographic History of Southern Chinese

**DOI:** 10.1101/2020.11.08.373225

**Authors:** Xiufeng Huang, Zi-Yang Xia, Xiaoyun Bin, Guanglin He, Jianxin Guo, Chaowen Lin, Lianfei Yin, Jing Zhao, Zhuofei Ma, Fuwei Ma, Yingxiang Li, Rong Hu, Lan-Hai Wei, Chuan-Chao Wang

## Abstract

Southern China is the birthplace of rice-cultivating agriculture, different language families, and human migrations that facilitated these cultural diffusions. The fine-scale demographic history *in situ*, however, remains unclear. To comprehensively cover the genetic diversity in East and Southeast Asia, we generated genome-wide SNP data from 211 present-day Southern Chinese and co-analyzed them with more than 1,200 ancient and modern genomes. We discover that the previously described ‘Southern East Asian’ or ‘Yangtze River Farmer’ lineage is monophyletic but not homogeneous, comprising four regionally differentiated sub-ancestries. These ancestries are respectively responsible for the transmission of Austronesian, Kra-Dai, Hmong-Mien, and Austroasiatic languages and their original homelands successively distributed from East to West in Southern China. Multiple phylogenetic analyses support that the earliest living branching among East Asian-related populations is First Americans (∼27,700 BP), followed by the pre-LGM differentiation between Northern and Southern East Asians (∼23,400 BP) and the pre-Neolithic split between Coastal and Inland Southern East Asians (∼16,400 BP). In North China, distinct coastal and inland routes of south-to-north gene flow had established by the Holocene, and further migration and admixture formed the genetic profile of Sinitic speakers by ∼4,000 BP. Four subsequent massive migrations finalized the complete genetic structure of present-day Southern Chinese. First, a southward Sinitic migration and the admixture with Kra-Dai speakers formed the ‘Sinitic Cline’. Second, a bi-directional admixture between Hmong-Mien and Kra-Dai speakers gave rise to the ‘Hmong-Mien Cline’ in the interior of South China between ∼2,000 and ∼1,000 BP. Third, a southwestward migration of Kra-Dai speakers in recent ∼2,000 years impacted the genetic profile for the majority of Mainland Southeast Asians. Finally, an admixture between Tibeto-Burman incomers and indigenous Austroasiatic speakers formed the Tibeto-Burman speakers in Southeast Asia by ∼2,000 BP.

## INTRODUCTION

Dated to ∼9,000 years before present (BP), Southern China is one of the two earliest agricultural centers in East Asia^1,2^. The First Farmers in Southern China domesticated a series of plants and animals indispensable for present-day people^3^, among which the most famous one is wet rice (*Oryza japonica*)^4^. It has long been hypothesized that the farming dispersal from Southern China involves human migration and in term the language families spoken by the farmers^5^. Recent studies have shown that the First Farmers in Southeast Asia^6,7^ and Pacific Islands^8-10^ derived most of their ancestry from a lineage shared with modern Southern Chinese, which has further been confirmed by ancient genomes from Fujian in coastal Southern China^11^ and adjacent Taiwan Island^12^. These finding supports the involvement of massive migration in the diffusion of Austronesian and Austroasiatic languages. However, our knowledge is still limited about the deep origin and early prehistory of such a ‘Southern East Asian’ or ‘Yangtze River Farmer’ lineage and its role in the genomic formation of modern Southern Chinese, especially the speakers of the other two indigenous language families in Southern East Asia, Hmong-Mien and Kra-Dai.

Here we generated new genome-wide data of 211 present-day Southern Chinese individuals, who belong to 30 geographic subgroups that have not yet been represented in genomic studies and cover the three main language families in this region, i.e., Hmong-Mien, Kra-Dai, and Sino-Tibetan. To thoroughly reconstruct the demographic history of Southern Chinese in relation to other East Asians, we co-analyzed them with ∼1,200 modern and high-coverage ancient samples from East and Southeast Asia, which cover the main ethnolinguistic and archaeological diversity in East Asia that is accessible till now with a high resolution.

Our study mainly addressed on three primary questions regarding the genomic history in East Asia. (1) How many ancestries substantially contributed to the gene pool of present-day East Asians and to what extent did they participate in the diffusion of different language families? Especially, if there are any sub-structure with the Southern East Asians? (2) What is the deep history regarding the origin of these ancestries and where did they originally inhabit? (3) What migrations and admixtures did these ancestries involve in the formation of present-day Southern Chinese? In response to questions above, we first gained an overview of genetic structure by principal component analysis (PCA)^13^ (Fig. 1A and B) and model-based clustering^14^ (Fig. 1C). To quantify the genetic affinity, we measured the degree of shared genetic drift^15^ (Fig. 2A) and shared haplotypes^16^ (Fig. 2B and C) among pairwise populations. We performed admixture graph modelling (Fig. 3A), admixture proportion estimation (Fig. 3B), and coalescent analysis based on site frequency spectrum (SFS) (Fig. 3C) to investigate the deep phylogenetic relationship and infer the geographic distribution of these ancestries. We then applied multiple methods, including the formal test of genetic homogeneity (Fig. 4), to further explore the admixtures in Southern China. Demographic history in East Asia and recent migrations and admixtures in Southern China has been summarized into map illustrations (Fig. 5). All the analyses above allow us to better decipher both the long-term prehistory and recent two millennia’s documented history along with Sinicization in Southern China from a genomic perspective.

**Figure 1.**
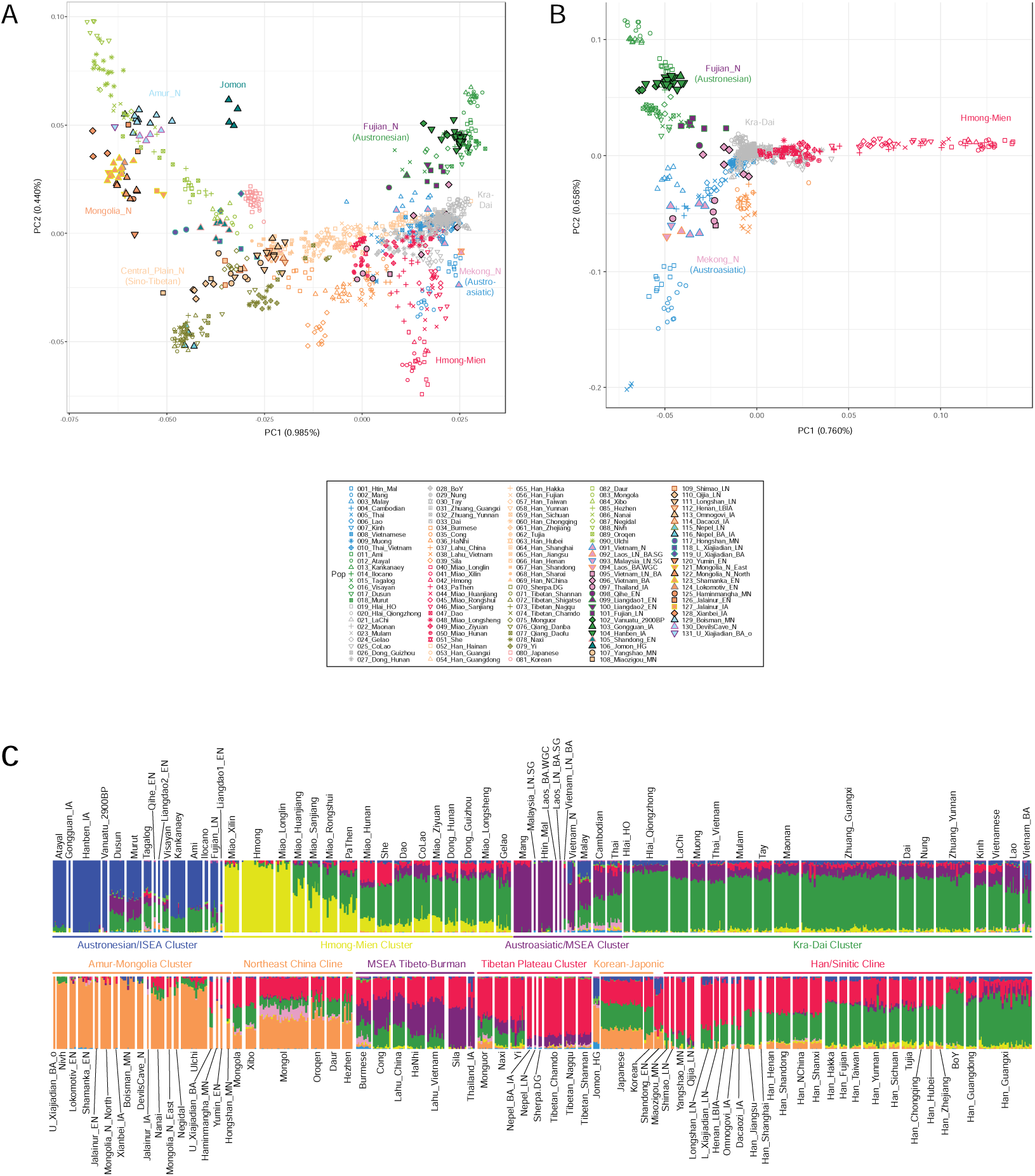
Genetic structure of East Asians. (**A to B**) PCA for (A) all the East Asians and (B) Southern East Asians. We projected ancient samples to principal components constructed by modern samples. (**C**) Unsupervised ADMIXTURE plot at K = 10, identifying six major ancestries in East Asia: orange, Northeast Asia/ Tungusic-related; red, Sino-Tibetan-related; blue, Austronesian-related; green, Kra-Dai-related; yellow, Hmong-Mien-related; purple, Austroasiatic-related.

**Figure 2.**
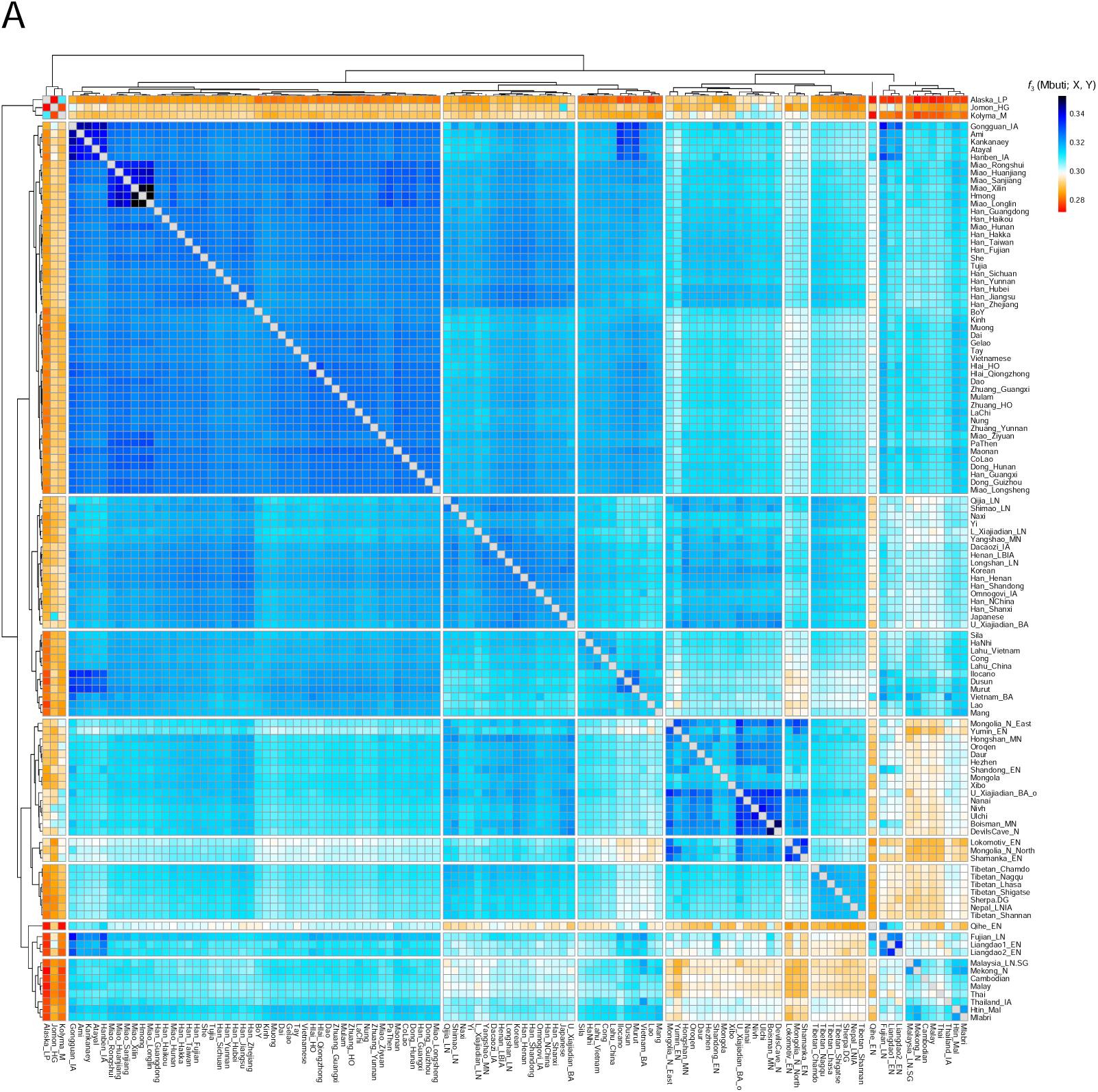
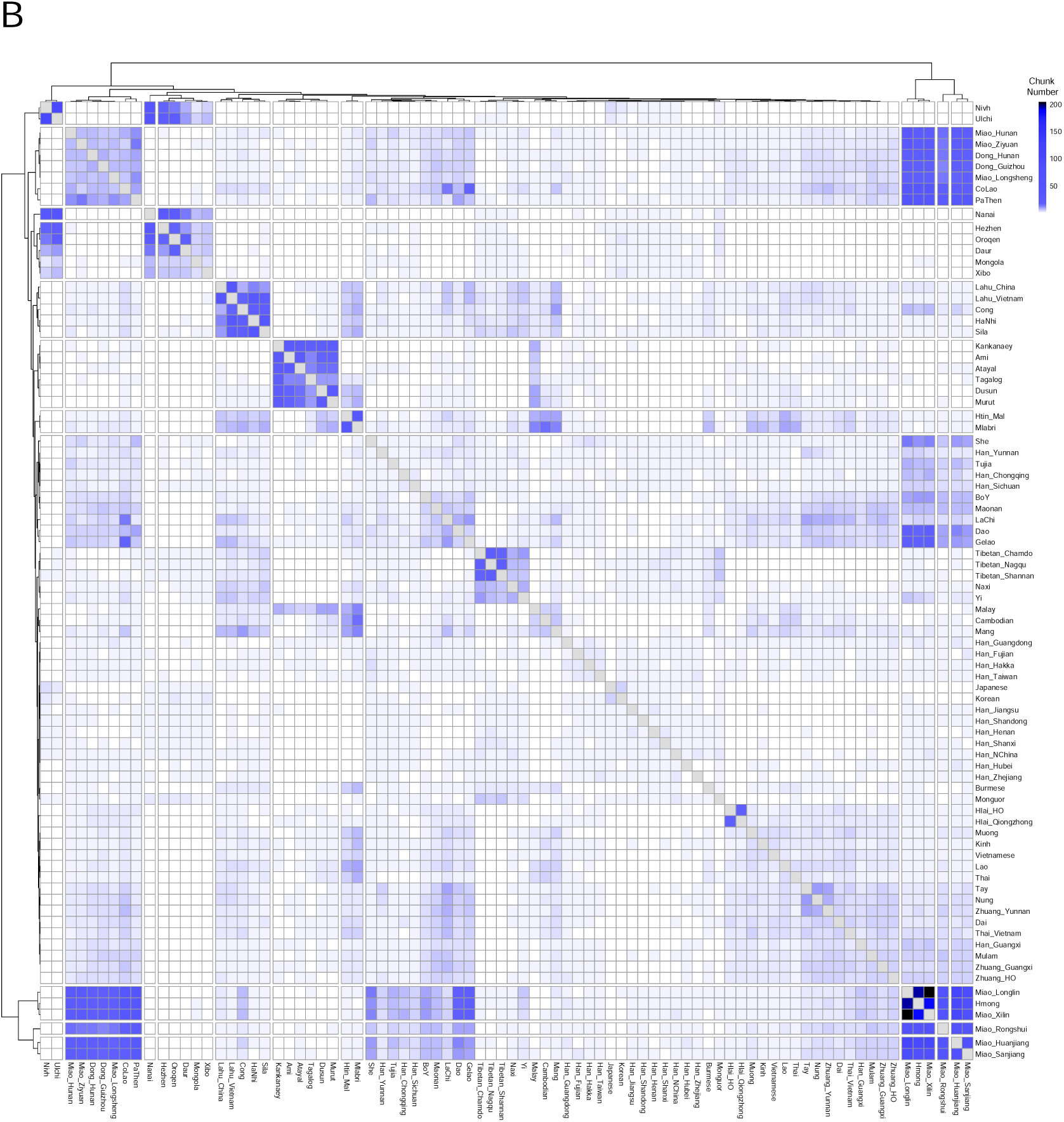
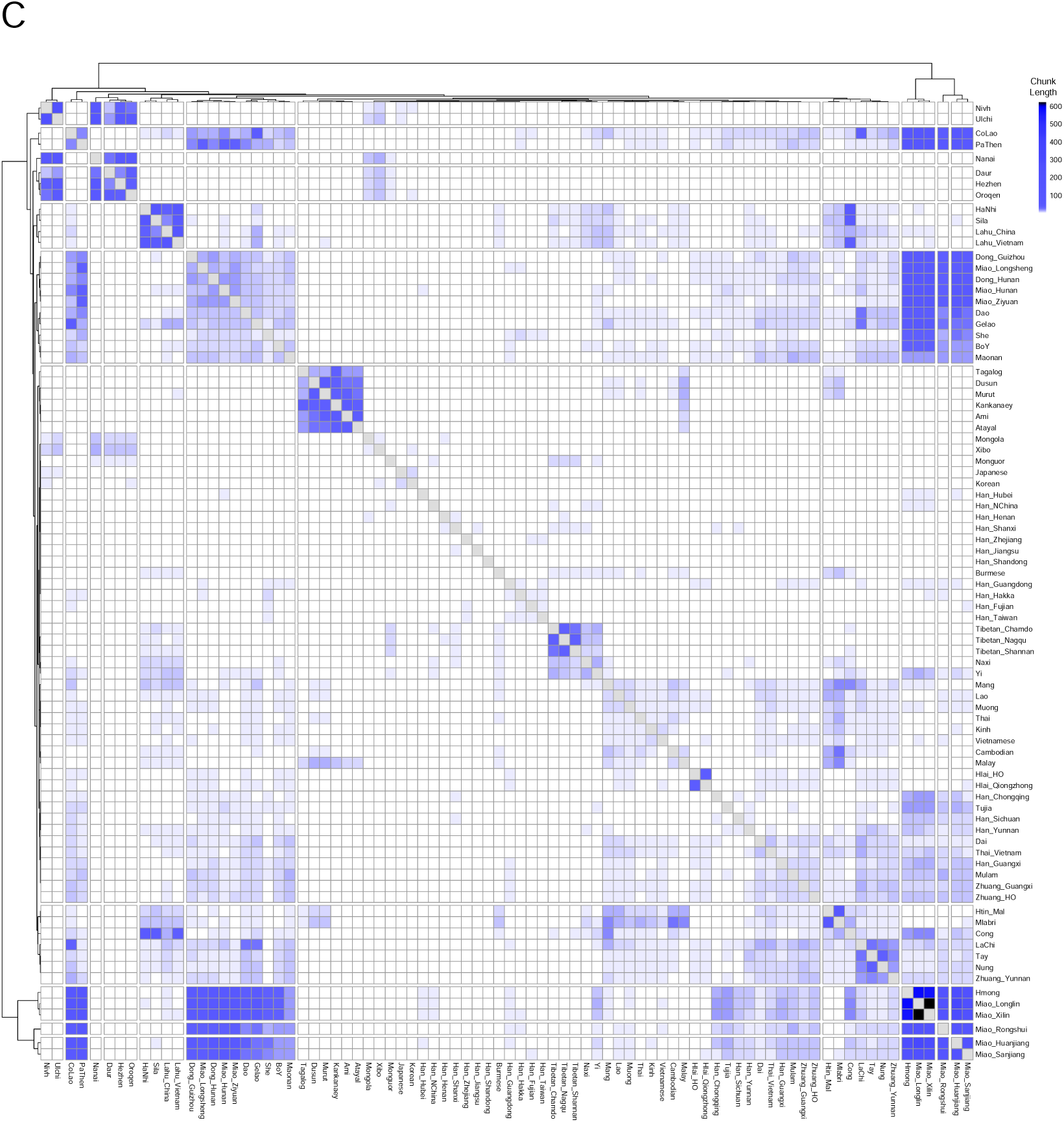
Quantitative measurement for pairwise genetic affinity. (**A**) Outgroup-*f*_*3*_ in the form *f*_*3*_(Mbuti; X, Y) measuring shared genetic drift between pairwise ancient and modern East Asian populations. (**B** to **C**) Normalized haplotype sharing based on (B) the number and (C) the total length (unit: cM) of shared IBD chunks for pairwise modern East Asian populations.

**Figure 3.**
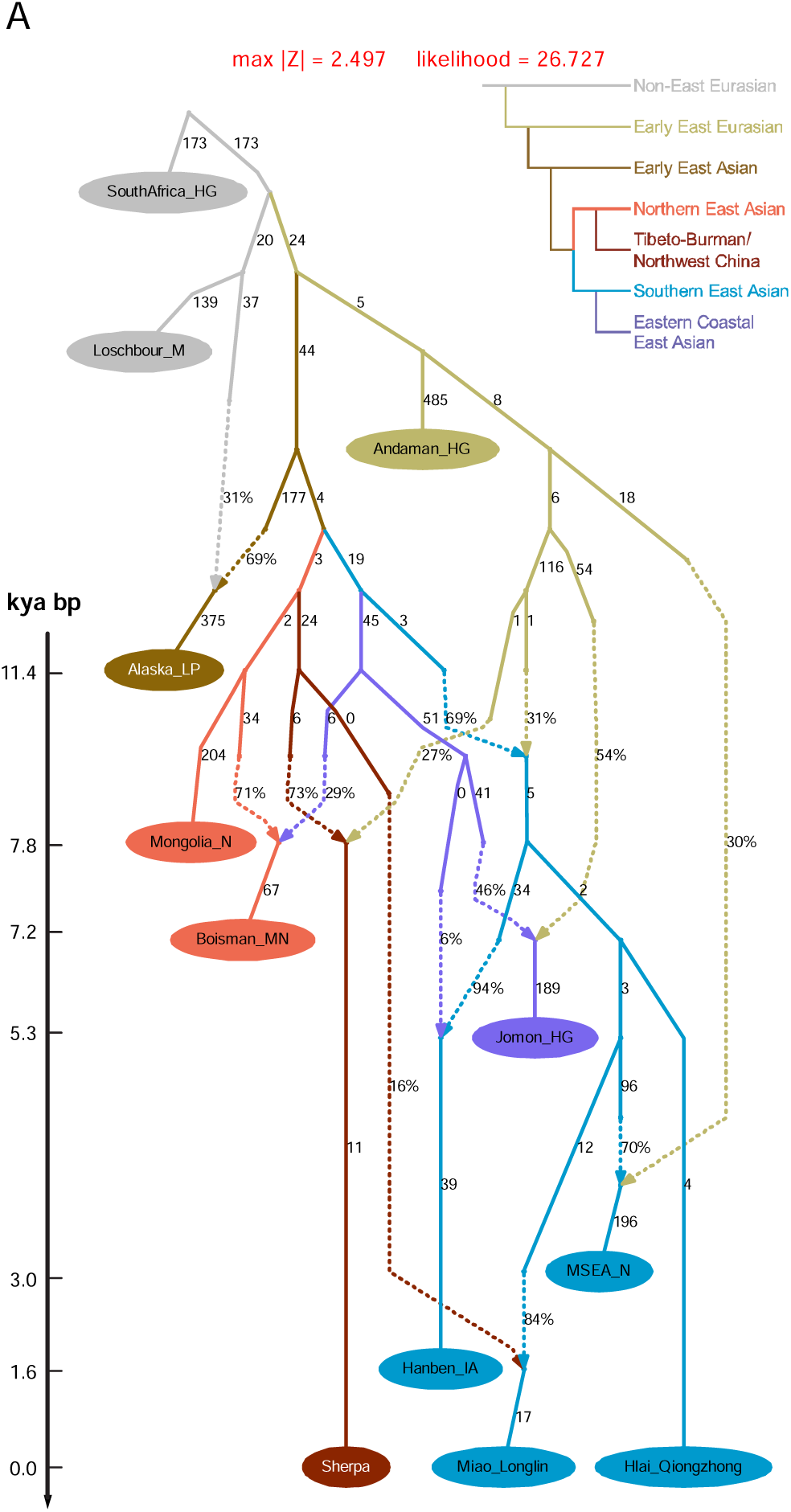
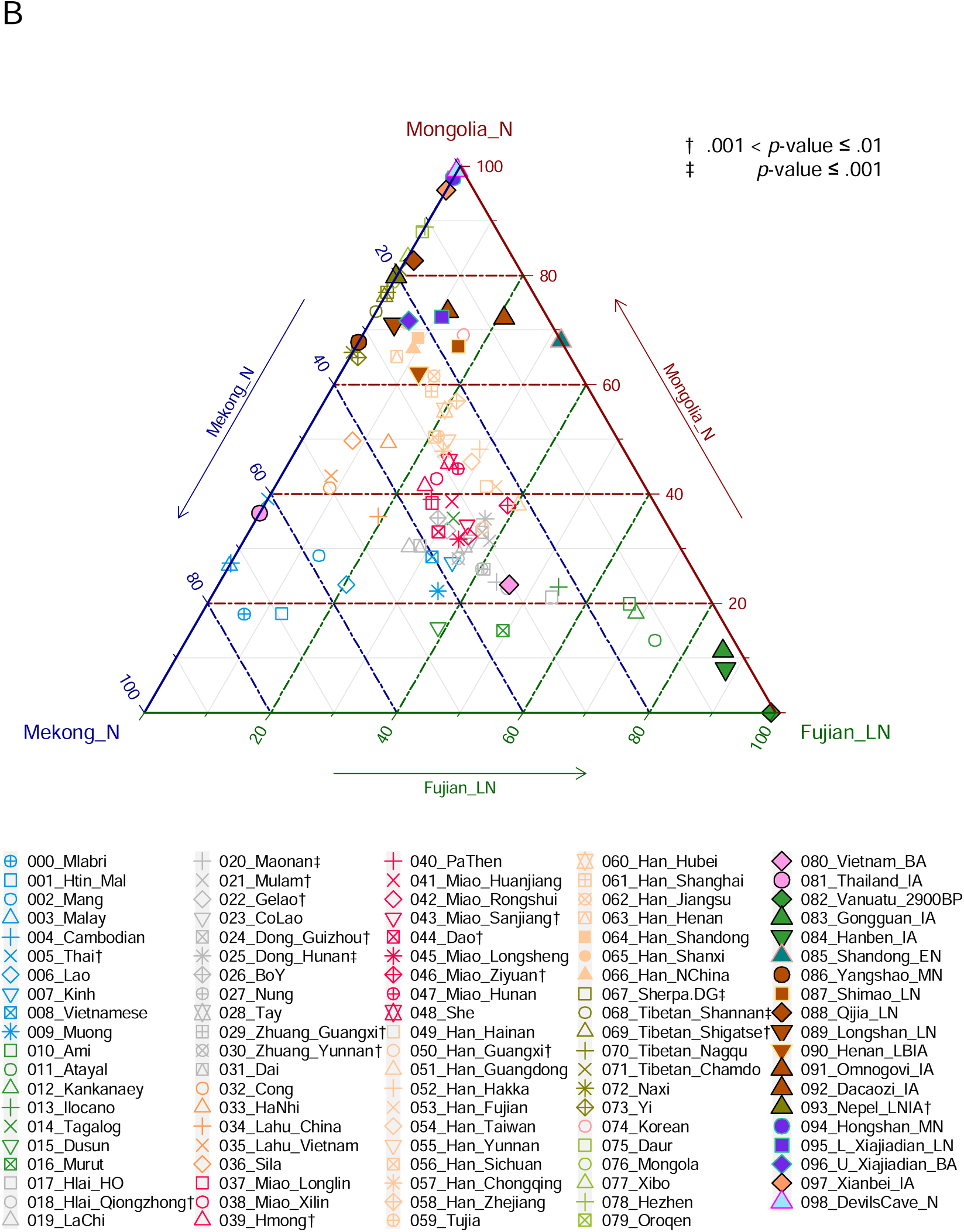
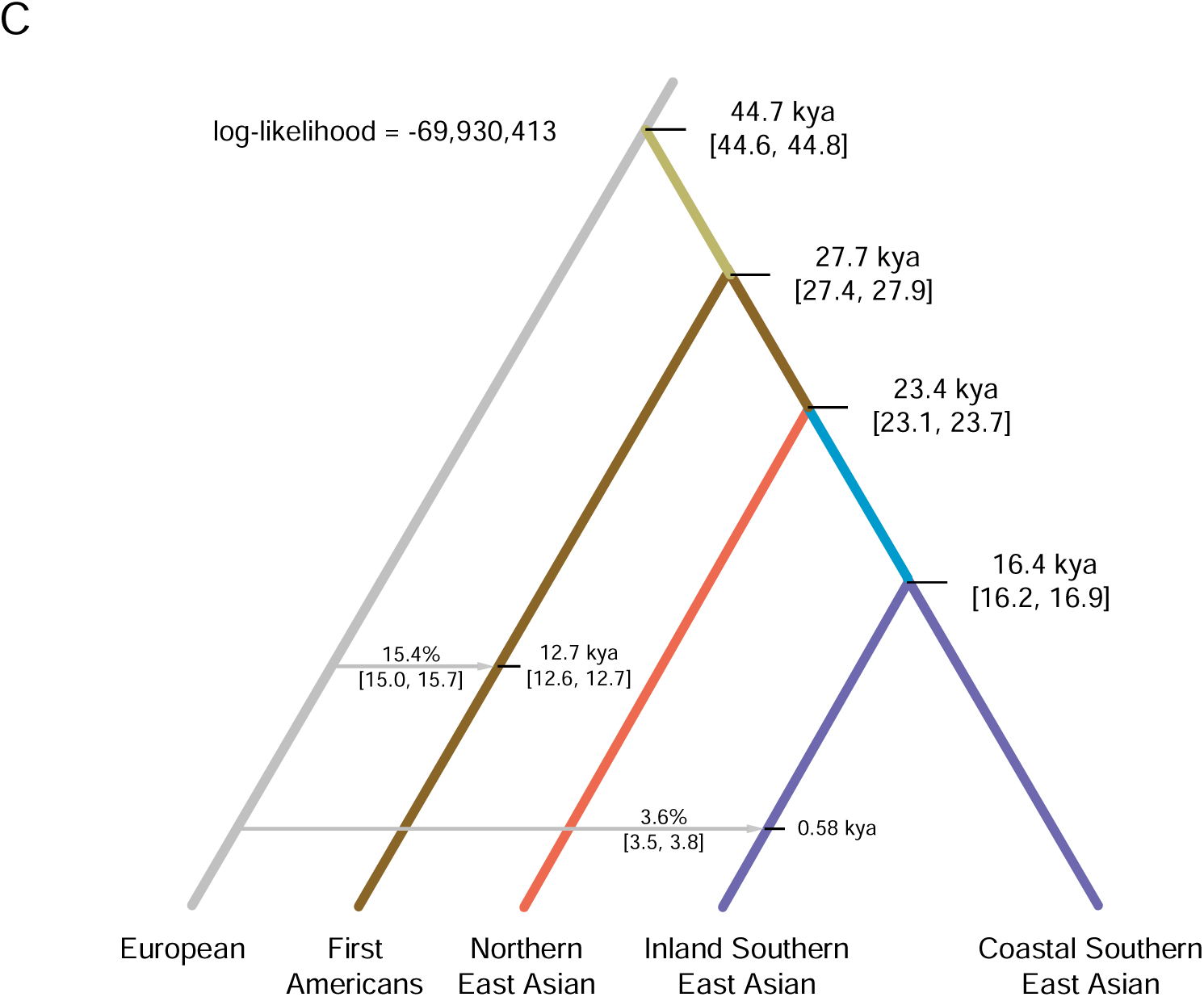
Demographic modelling for deep history of East Asians. **(A)** Optimal *qpGraph* admixture model for the phylogenetic relationship among the surrogates for major ancestries in East Asia. Alaska_LP, Mongolia_N/Boisman_MN, Sherpa, Hanben_IA, Hlai_Qiongzhong, Miao_Longlin, and MSEA_N respectively surrogate First Americans, Northeast Asian, Sino-Tibetan, Austronesian, Kra-Dai, Hmong-Mien, and Austroasiatic ancestries. Drift along each edge are multiplied by 1,000. (**B**) Three-source *qpAdm* models the contribution of Northern East Asian (represented by Mongolia_N), Coastal Southern East Asian (represented by Fujian_LN), and Inland Southern East Asian (represented by Mekong_N) lineages in ancient and present-day East Asians. (**C**) Coalescent analysis using SFS of rare alleles to calibrate the time of the major splits in East Asians (implemented by Rarecoal). We used whole genome sequences from 56 individuals in this analysis. kya, thousand years ago.

**Figure 4.**
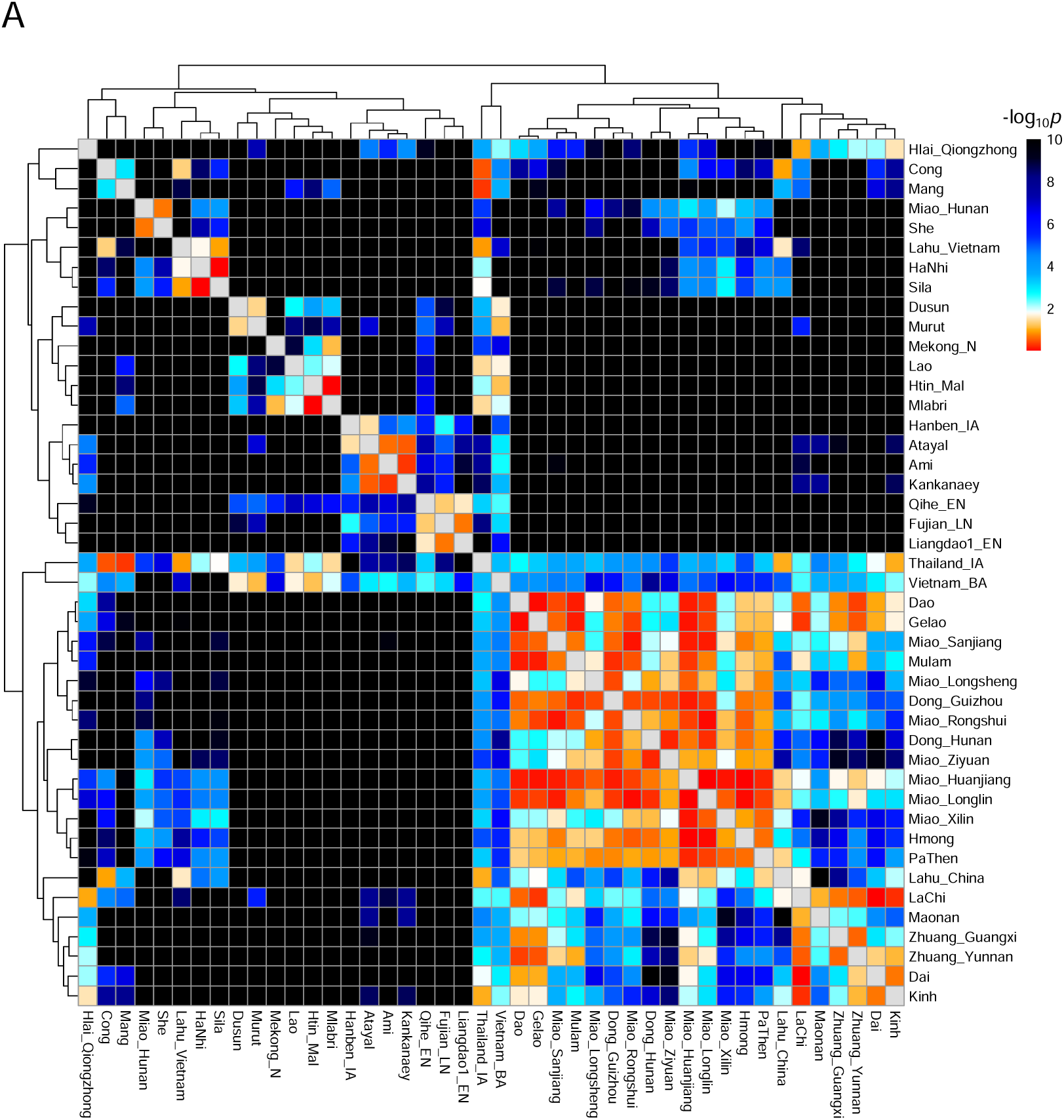
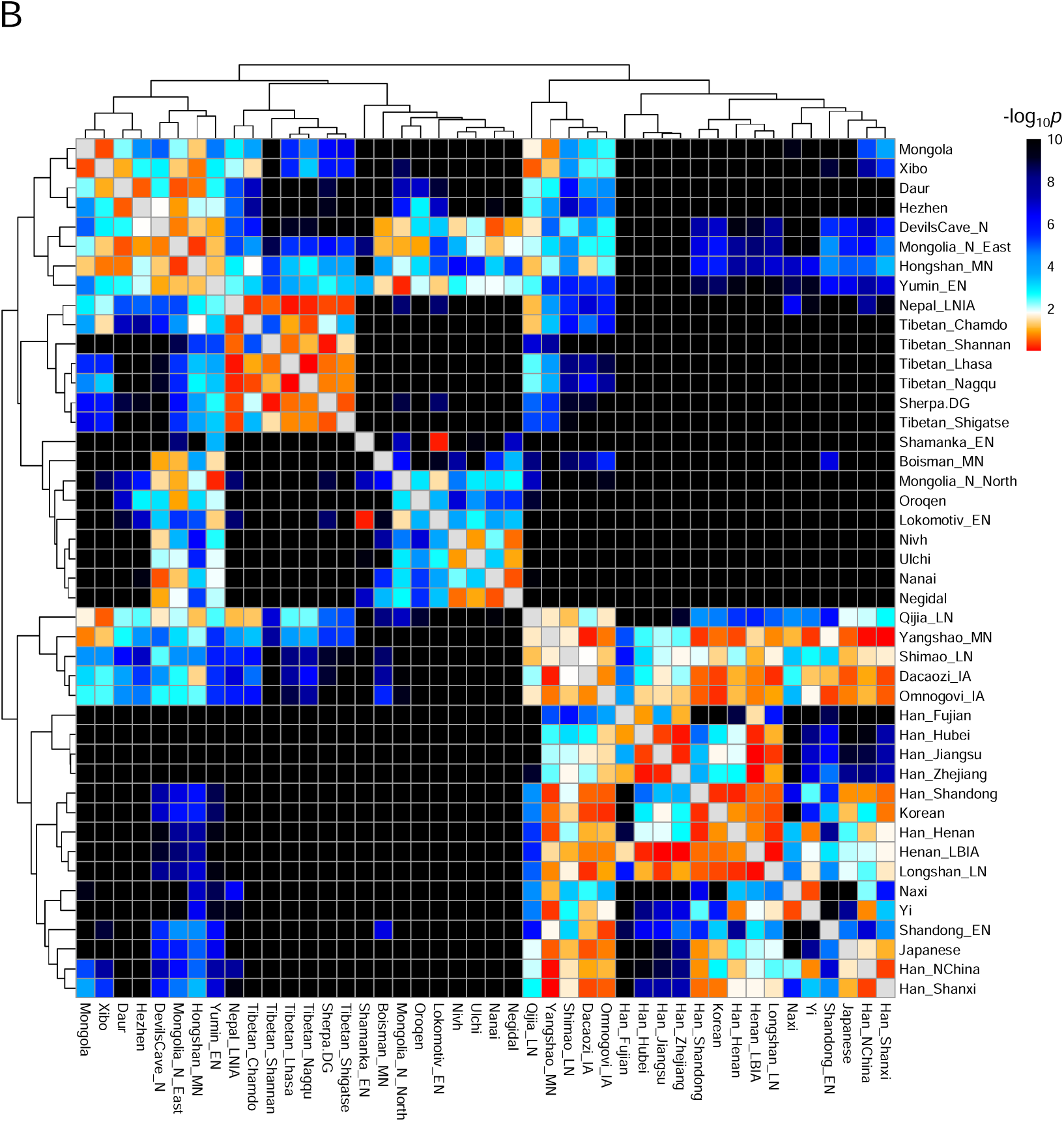
Genetic homogeneity of pairwise populations. Heatmaps show negative logarithms for *p*-values of pairwise *qpWave* in (A) Southern East Asians and (B) Northern East Asians.

**Figure 5.**
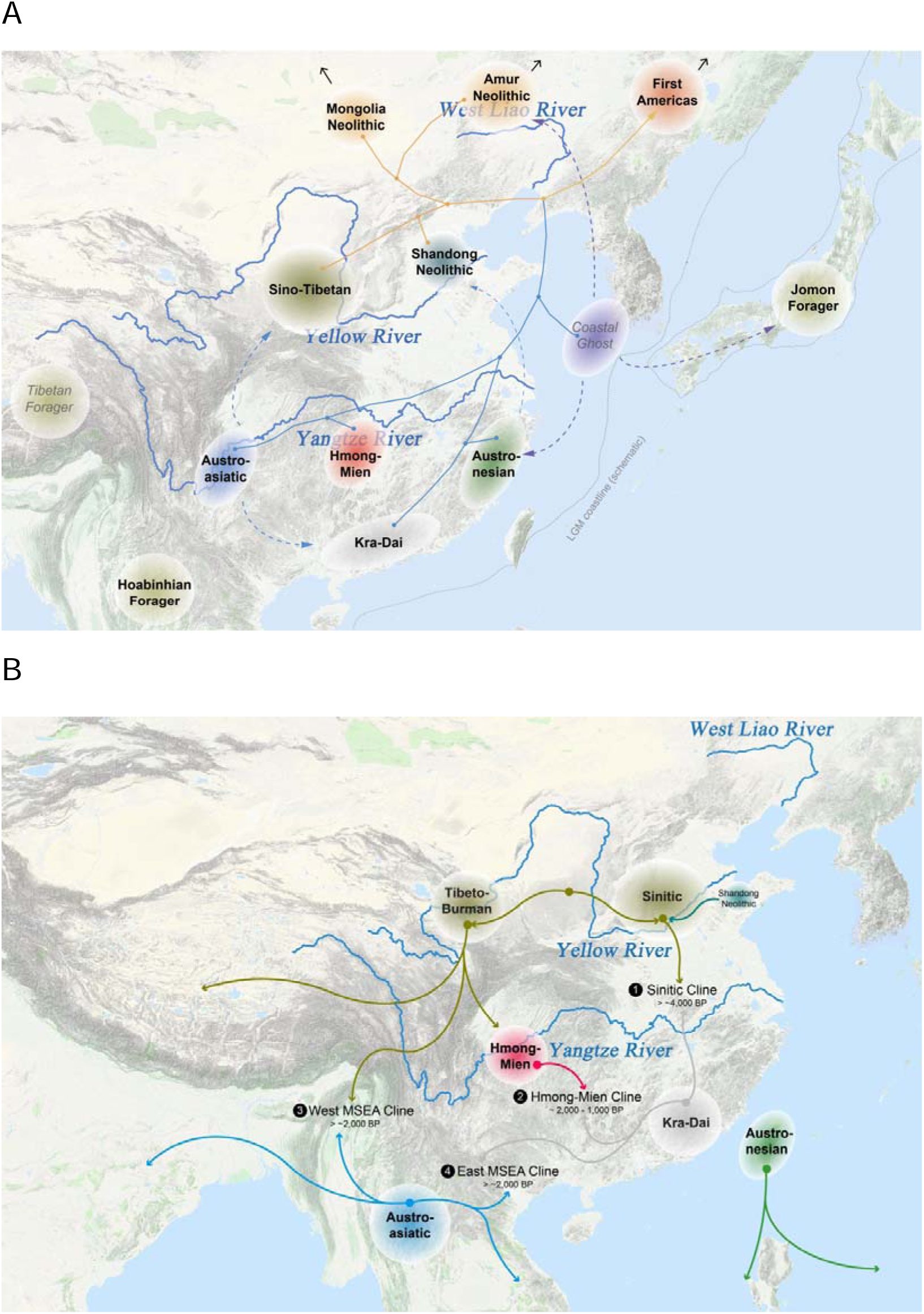
Illustrations for demographic history in East Asia. (**A**) The formation of geographically and linguistically structured ancestries in East Asia. (**B**) Massive migrations and admixtures forming the current genomic landscape in Southern China and neighboring regions.

## RESULTS

### A. Genetic structure in East Asia highly corresponds to linguistic affiliations

PCA of all the ancient and modern East Asians in our dataset (Fig. 1A) replicates the pattern of genetic differentiation between Northern and Southern East Asians^11,12^ and variations between two geographically structured clusters within Northern East Asia^12,17^. The ‘Inland Yellow River Cluster’ is best represented by Neolithic Farmers from Inland Yellow River Basin (Yangshao_MN and Qijia_LN^17^) and later and modern Tibetan Plateau inhabitants. In ADMIXTURE plot (Fig. 1C), the ancestry maximizing in this cluster is also ubiquitous in modern Sino-Tibetan speakers (Fig. 1C), consistent with previous linguistic studies that suggested a plausible origin of Proto-Sino-Tibetan in Yangshao Culture^18,19^. The ‘Northeast Asian Cluster’ is typified by the Neolithic hunter-gatherers in Amur River Basin (DevilsCave_N^18,19^ and Boisman_MN^12^), Mongolian Plateau, and Cis-Baikal (Shamanka_EN and Lokomotiv_EN^19^), as well as modern populations from Amur River region (Fig. 1A), and these surrogates also harbor the highest proportion of the ancestry prevalent in modern Tungusic speakers and other Northeast Asians. We observe consistent clustering manners in pairwise outgroup-*f*_*3*_ (Fig. 2A) and haplotype sharing degree measured by identity-by-descent (IBD) chunks (Fig. 2B and C).

Confirming the highly shared genetic history among the Southern East Asians, we find that Kra-Dai and Hmong-Mien populations in Southern China, Southern Han Chinese, and Austronesian Taiwanese cluster together with a high sharing degree (*f*_*3*_ > 0.320) in outgroup-*f*_*3*_ matrix (Fig. 2A). We further identify a previously unknown genetic structure within the Southern East Asians that has a strong connection with linguistic classification. That is, each of the speakers of all the four indigenous language families in Southern China and Southeast Asia has their own cluster in PCA for Southern East Asians (Fig. 1B) and their own ancestral component in ADMIXTURE plot (Fig. 1C). Austronesian-related ancestry maximizes in ancient and modern Austronesian Taiwanese with a high proportion in earlier Fujian Neolithic individuals. Austroasiatic-related ancestry maximizes in Neolithic farmers in Mekong River Basin (collectively referred as ‘Mekong_N’ afterwards) and is also extensively distributed in Mainland Southeast Asia. Hmong-Mien-related ancestry maximizes in the Western Hmong populations (Miao_Longlin and Miao_Xilin) newly reported in this study and is absent in all the ancient individuals, indicating that the original homeland of this ancestry has not yet been sampled in ancient genomic studies to date. Kra-Dai-related ancestry maximizes in modern Kra-Dai speakers resident in Mainland Southern China (Zhuang, Maonan, Mulam) and neighboring Hainan Island (Hlai). Especially in Austronesian- and Hmong-Mien-speaking populations, we observe a high degree of IBD sharing between the surrogate populations with other populations of the same language affiliation [number > 11.5, length > 18.5 cM for Atayal; number > 12.4, length > 32.0 cM for Miao_Longlin], which indicates strong genetic drift or founder effect in both ancestries. We use the terms ‘Core Austronesians/ Kra-Dais/ Hmong-Miens/ Austroasiatics’ to address modern surrogates of each of the ancestries.

We implemented point-biserial correlation to formally examine if the proportion of ancestries in ADMIXTURE is significantly associated with corresponding language families (Extended Data Table 2), which shows a strong correlation to linguistic affiliation for all the six major ancestries in East Asia (r_pb_ > 0. 500, *p* < 0.0001). The gene-language correlation indicates that present-day speakers of the major language families in East Asia usually receive a strong genomic legacy from the speakers of corresponding proto-languages, which enables us to trace the history of these language families through the history of human populations.

### B. Reconstruction of ancestral homelands in Southern China

One of the crucial historical aspects for a language family is its original homeland, or *Urheimat*. In historical linguistics, two main strategies have been applied for the reconstruction of linguistic homeland^20^. One of the strategies leverages the reconstructed vocabulary of the proto-language to inform us about the environment and lifestyle of the original speakers, which is in turn a clue for geographic origin. The other strategy assumes that the linguistic homeland tends to be located in where there is a higher linguistic diversity^21^.

However, both linguistic strategies cannot work well in case of the four major language families that are indigenous in Southern East Asia and are supposed to originate from Southern China. First, all of the four language families share some words reflecting either a common origin or extensive language contact among all of them, especially the ones related to farming and domestication^22^, which prevents us to infer the homeland distribution within Southern China in a higher resolution. Second, massive cultural transition happened in Southern China in recent millennia, especially Sinicization^23^, may have largely decreased language diversity in this region. Given the language-associated genomic structure in Southern East Asia, here we adapted another genomics-based strategy — modelling the phylogenetic relationship among the ancestries to infer their relative geographic distribution, hence the homelands for the language families they spread.

We implemented *qpGraph*^24^ to model the phylogenetic relationship of the main ancestries in East Asia, using populations in which the corresponding ancestries maximize in PCA (Fig. 1A and B) and ADMIXTURE (Fig. 1C) as surrogates. Our optimal model (Fig. 3A) shows that Northern East Asians (Mongolia_N, Boisman_MN, and Sherpa) and Southern East Asians (Hanben_IA, Hlai_Qiongzhong, Miao_Longlin, and MSEA_N) each form a monophyletic lineage, with additional admixture from an Andamanese-related deep ancestry in the common ancestor of Southern East Asians (31%). As Core Kra-Dais, Core Hmong-Miens, and the earliest Austronesian-related ancient group (Qihe_EN and Liangdao_EN) are located in Southern China, we can reasonably deduct that the common ancestor of Southern East Asian lineage and its initial diversification also took place in Southern China. Sub-topology within the Southern East Asian lineage suggests that Austronesian ancestry split with the others first, then followed by Kra-Dai ancestry, with Hmong-Mien and Austroasiatic ancestries sharing the most genetic drift with each other. Given the easternmost and westernmost geographic position respectively for the earliest Austronesian- (Qihe_EN and Liangdao_EN) and Austroasiatic-related (Mekong_N) individuals, such a topology can be explain as a result of the isolation-by-distance pattern from east to west in ancient Southern China sequentially for Austronesian, Kra-Dai, Hmong-Mien, and Austroasiatic ancestries (Fig. 5A).

To test if the pattern observed above is more extensively applicable in East Asian populations, we then used *qpAdm*^25^ to parse the contribution of Northern, Coastal Southern, and Inland Southern East Asian lineages, respective represented by Mongolia_N (as they have minimal Southern East Asian ancestry in admixture graph), Fujian_LN, and Mekong_N (Fig. 3B). Core Austronesians and Core Austroasiatics respective derive most of their ancestry related to Fujian_LN (66.9%–74.3%) and Mekong_N (58.0%–75.2%), suggesting that the Neolithic genomic structure in Southern East Asians has largely preserved in these modern isolated populations. In present-day Southern China, Core Kra-Dais harbors more of their ancestry related to Fujian_LN (39.0%–53.9%) than Mekong_N (24.9%–32.3%) and we also observe a similar pattern in Southeast Han Chinese (28.9%–40.3% for Fujian_LN and 21.8%–25.2% for Mekong_N), suggesting that the Kra-Dai ancestry itself primarily derives from an Austronesian-related lineage with additional Austroasiatic-related gene flow. On the contrary, Core Hmong-Mien derives more of their ancestry related to Mekong_N (32.3%–35.3%) than Fujian_LN (23.7%–26.4%), consistent with the closest phylogenetic relationship of Austroasiatic and Hmong-Mien ancestries in admixture graph (Fig. 3A). Regardless different topology between Kra-Dai and Austronesian ancestries suggested by *qpGraph* and *qpAdm*, both analyses indicate a consistent pattern for original geographical distribution of these four ancestries (Fig. 5A).

We used admixture-*f*_*3*_ statistics^24^ to examine two alternative explanations for phylogenetic relationship between Kra-Dai and Austronesian ancestries. Due to small effective population size (N_E_) and strong genetic drift in many East Asian populations (Extended Table Fig. 1), we are usually unable to obtain significant negative *f*_*3*_. Therefore, we focused on the lowest *f*_*3*_ value given by different pairs of source populations and exhausted all the potential pairs. The results might not be statistically significant, but it can provide a consistent pattern if the same pair of source populations minimizes *f*_*3*_ of a series of different target populations. For most of the Kra-Dai-speaking populations and Vietic speakers who have a genetic profile resembling their Kra-Dai neighbors (Extended Table 3C), the lowest *f*_*3*_ achieves when one of the source population surrogates Austroasiatic ancestry (Mekong_N, Malaysia_LN, and Mlabri) and the other surrogates Austronesian ancestry (Fujian_LN and Austronesian Taiwanese), with the strongest negative signal in Kra-Dai and Vietic speakers in Mainland Southeast Asia (Muong, Nung, Lao). Since all of these populations live in the northern interior of Mainland Southeast Asia, a direct and recent gene flow from Austronesian speakers does not seem to be a feasible scenario. A more possible scenario is that the Kra-Dai ancestry itself, at least partially, is closely related to Austronesian ancestry and this favors the model suggested by *qpAdm*. The partial Austronesian-related origin for Kra-Dai ancestry is compatible with the ‘Austro-Tai’ hypothesis in historical linguistic^26-28^ that suggests a common origin of Austronesian and Kra-Dai language families, and the motif Y-chromosomal haplogroup O1a-M119 that is dominant in Neolithic Fujian individuals^11^ and shared by Austronesians, Kra-Dais, and Southern Han Chinese^29^.

The impact of distinct Southern East Asian ancestries is not limited within Southern China and further south. In the earliest samples from Yellow River Basin, coastal individuals from Early Neolithic Shandong derive all of the southern ancestry from a Fujian_LN related lineage (32.0 ± 6.6%, Fig. 3B), consistent with their Austronesian-related ancestral component in ADMIXTURE (Fig. 1C) that largely disappears in later Northern East Asians. In contrast, inland individuals from Middle Neolithic Yangshao Culture derive all of the southern ancestry from a Mekong_N related lineage (32.2 ± 5.9%, Fig. 3B). These genomes document that the initial isolation and genomic differentiation among geographically structured populations in Southern China is no late than Early Neolithic and demographic contact between Northern and Southern China in this period is via distinct coastal and inland routes (Fig. 5A).

Instead of a direct contribution from contemporary Neolithic Fujian population, a more plausible scenario for the Austronesian-related ancestry in Shandong Neolithic individuals they received a gene flow from a currently unsampled ancient population from neighboring regions like Jiangsu or Zhejiang, who in turn harbored a Fujian_LN-related lineage (Fig. 5A). This scenario is also consistent with the strong connection between Neolithic cultures in Lower Yangtze Basin, like Hemudu Culture, and contemporary Fujian, like Tanshishan Culture, to which the Fujian_LN individuals belong^11^. Taking genomic, linguistic, and archaeological evidence together into account, Kra-Dai ancestry likely originates from the local Austronesian-related lineage in continental Southeast China approximately ranging from Zhejiang to Guangdong with additional gene flow from an Austroasiatic-related lineage (Fig. 5A), whereas the migrants from continental Southeast China to Taiwan largely preserve the original Austronesian/Austro-Tai-related genetic profile and are responsible for the massive Austronesian expansion. However, the coexistence of Kra-Dai and Austronesian ancestry once in the continent still cannot be fully excluded in light of our analysis.

With nearly absolute southern affinity to Austroasiatic-related lineage (Fig. 3B), the genetic profile of Yangshao individuals largely persists in Late Neolithic individuals from Qijia Culture (16.0 ± 8.3% for Mekong_N, 1.3 ± 10.6% for Fujian_LN) –whose expansion is supposed to parallel with the diffusion of at least some of the Tibeto-Burman languages^30^–and modern Tibeto-Burman speakers in Tibetan Plateau (18.9–24.2% for Mekong_N, 0.0% for Fujian_LN) and Tibetan-Yi Corridor like Naxi and Yi (33.7–34.1% for Mekong_N, 0.0–1.3% for Fujian_LN). In admixture-*f*_*3*_ (Extended Data Table 3A), Tibeto-Burman populations in Tibetan-Yi Corridor and further south show a consistent pattern of two-way admixture by Sino-Tibetan and Austroasiatic ancestries. Multiple evidences suggest that populations with Austroasiatic ancestry likely distributed further north in Southwest China previously. Given the close relationship between Austroasiatic and Hmong-Mien ancestries, it is reasonable to deduct that the place of origin for both ancestries is in Southwest China (Fig. 5A) and both of them are possibly related to the Neolithic farming cultures in Middle Yangtze, e.g., Daxi Culture^3^, which is also consistent with that modern populations with significant Hmong-Mien-related ancestry (Fig. 1C) are distributed in Guangxi, Guizhou, and Hunan of Southwest China.

### C. Deep history of East Asian populations

Genetic drift-based admixture graph analysis by *qpGraph* is informative and robust for phylogenetic reconstruction with admixture events, but it cannot estimate the time of splits and admixtures since genetic drift is not proportional to time^24^. Therefore, we obtained coalescent time estimation using SFS-based framework Rarecoal^31^ and whole genome sequences from Europeans, First Americans (Mixe, Piapoco, and Pima), Northern East Asians (Ulchi, Hezhen, and Oroqen), Coastal Southern East Asians (Ami, Atayal, and Igorot), and Inland Southern East Asians (Cambodian and Thai). We estimated that the divergence between East and West Eurasians is ∼44,700 BP (95% confidence interval (CI) 44,600–44,800 BP, Fig. 3C), consistent with the time estimation in previous studies^19,32^ and the equal genomic relationship to East and West Eurasians for the ∼45,000-year-old Ust’-Ishim individual^33^.

Earlier works have discovered that the First Americans primarily derive from an East Asian-related lineage with additional admixture with West Eurasian-related Ancient North Siberians^15,19^. However, it is still unclear how the East Asian-related ancestry of First Americans relates to other East Asians. Both *qpGraph* model (Fig. 3A) and Rarecoal model (Fig. 3C) suggest that the East Asian ancestor of First Americans represents the deepest East Asian-related lineage in all the living population who split with the common ancestor of both Northern and Southern East Asian lineages ∼27,700 BP (95% CI 27,400–27,900 BP, Fig. 3C). After that, the ancestor of Northern and Southern East Asians split with each other ∼23,400 BP (95% CI 23,100–23,700 BP), which is prior to the Last Glacial Maximum (LGM) in East Asia (∼21,000–15,000 BP)^3^. This implies that the differentiation between Northern and Southern East Asian lineages plausibly results from the isolation of geographically structured populations in different refugia during LGM. Within Southern East Asian lineage, the separation between Coastal and Inland Southern East Asian lineages took place ∼16,400 BP (95% CI 16,200–16,900 BP), which is contemporary with LGM and significantly earlier than the earliest farming practice in Southern China (∼9,000 BP)^3^. Such a result indicates that the Neolithic transition for different Southern East Asian ancestries might result from either independent acquisition or the spread of idea without massive population replacement.

The genomic origin of Jomon hunter-gatherers in Japanese Archipelago is mysterious due to their basal East Eurasian ancestry compared with other East Asians and their additional genetic affinity to Amur Basin populations and Austronesian Taiwanese^6,12,34^. In our admixture graph model (Fig. 3A), there are two different layers contributing to the genetic profile of Jomon hunter-gatherers. The first layer is distantly related to Andamanese hunter-gatherers, which is likely introduced by an earlier peopling of Japanese Archipelago. The second layer is a sister lineage of Southern East Asian, which explains the genetic affinity of Jomon to other coastal East Asian populations. Compared with the large proportion of Andaman-related ancestry in Jomon hunter-gatherers (56.5 ± 4.8%), the small amount of Andamanese-related ancestry in ancient (6.4–11.7%) and modern (1.0–2.1%) populations in Amur Basin also suggests that their affinity to Jomon hunter-gatherers is more feasible to be explained by an East Asian-related ancestry than an Andamanese related ancestry. Taking both *qpGraph* (Fig. 3A) and Rarecoal (Fig. 3C) into account, the formation of this sister lineage to Southern East Asian is between ∼23,400 BP and ∼16,400 BP, which mostly falls in the range of LGM. Therefore, a plausible geographic distribution for this lineage is in the continental coastal East Asia, which is largely below the sea level at present (Fig. 5A).

### D. Migrations and admixtures shaping present-day Southern Chinese

East Asia in recent millennia has witnessed a series of massive demographic events that contribute to the formation of modern East Asians. Here we particularly focus on Southern China and characterize the most crucial migrations and admixtures revealed in light of our results (Fig. 5B).

#### (D.1) Formation of Han Chinese

Han Chinese comprise around one fifth of the world’s population^35^. Previous studies suggest that Han Chinese is primarily formed by Yellow River Farmers (i.e., Sino-Tibetan ancestry in this study) with additional gene flow from Southern East Asian lineage^12^. However, it is still not fully known which specific ancestry mostly contribute to the southern ancestry of Han Chinese. In ADMIXTURE plot (Fig. 1C), both Northern and Southern Han Chinese have a similar genetic profile comprising both Sino-Tibetan and Kra-Dai ancestries, with an increase of Kra-Dai ancestry from North to South. The earliest individuals with such a genetic profile in Yellow River Basin are from Longshan Culture (∼4,000 BP) and the genetic profile in individuals resembling the genetic profile of Northern Han Chinese is found in Dacaozi_IA and Omnogovi_IA (previously assigned as Xiongnu individuals^36^) during Han Dynasty (∼4,000 BP). Formal test for pairwise genetic homogeneity conducted by *qpWave* (Fig. 4B) confirms that the genetic homogeneity between Han Dynasty individuals and any of Neolithic Shandong individuals and Inland Yellow River individuals (Yangshao_MN and Qijia_LN) is higher than the one between the latter two, consistent with a closer position for Han Dynasty individuals than Neolithic Yellow River individuals in PCA (Fig. 1A) and *qpAdm* (Fig. 3B). This indicates that the admixture between Inland and Coastal Yellow River plays an important role in the formation of Northern Han Chinese. Regarding the formation of Sinitic Cline and Southern Han Chinese, admixture-*f*_*3*_ results (Extended Table 3B) suggest that the strongest signal of admixture come from the pair of surrogates for Sino-Tibetan ancestry (Qijia_LN) and Kra-Dai (Hlai) or Austronesian ancestry (Ami, Atayal, and Kankanaey). Therefore, we conclude that the Sinitic Cline is primarily formed by massive southward migration of Northern Han Chinese and subsequent admixture with indigenous Kra-Dai speakers in Southern Chinese.

#### (D.2) Admixture between Hmong-Mien and Kra-Dai populations

Another major genetic cline in South China is the Hmong-Mien Cline, which comprises most of the Hmong-Mien speakers as well as neighboring Kra-Dai populations in the interior of Southern China (Fig. 5B). In ADMIXTURE plot (Fig. 1C), Hmong-Mien populations from west to east show a decrease of Hmong-Mien ancestry and an increase of Kra-Dai ancestry, suggesting that the migration of Kra-Dai speakers came from the east and constantly admixed with local Hmong-Mien populations. Meanwhile, adjacent Kra-Dai speakers of Kra (Gelao) and Kam-Sui (Dong) branches also receive significant Hmong-Mien ancestry, indicating a bidirectional gene flow. Admixture time estimation performed by ALDER (Extended Data Table 4) shows that the admixture of Hmong-Mien Cline happened ∼24–46 generations ago overlapping Tang Dynasty to Yuan Dynasty.

#### (D.3) Spread of Kra-Dai ancestry in Mainland Southeast Asia

Besides the contribution to Han Chinese and Hmong-Mien populations, Kra-Dai ancestry also has a strong impact to Mainland Southeast Asia in recent two millennia. The earliest genomic document for the arrival of Kra-Dai ancestry in Mainland Southeast Asia is Bronze Age individuals from northern Vietnam (∼2,000 BP, Fig. 1C). Further extensive admixture between populations with more Kra-Dai ancestry and more Austroasiatic ancestry that arrived earlier largely explain the more obvious discrepancy between gene and language in Mainland Southeast Asians than other Southern East Asians. Especially, it is evident in PCA (Fig. 1A and B), ADMIXTURE (Fig. 1C), and outgroup-*f*_*3*_ (Fig. 2A) that Austroasiatic-speaking Kinh and Muong of the Vietic branch have a more similar genetic profile with Kra-Dai speakers in South China than other Austroasiatic speakers with a more typical ‘Austroasiatic’ genetic profile, suggesting a language shift from incoming Kra-Dai language to local Austroasiatic language as a possible mechanism.

#### (D.4) Genomic origin of Tibeto-Burman-speaking Mainland Southeast Asians

The special Y-chromosomal haplogroup F2-M427 in Lahu^37^ and many other Tibeto-Burman populations in Mainland Southeast Asia raise further question for their genomic origin. In ADMIXTURE plot (Fig. 1C), we find that Tibeto-Burman speakers in Mainland Southeast Asia (Lahu from China and Vietnam, Sila, HaNhi (Hani), and Cong) majorly have a genetic profile comprising Sino-Tibetan and Austroasiatic ancestries, with a consistent pattern in *qpAdm* (36.7–50.1 % for Mekong_N, 7.9–19.1% for Fujian_LN, Fig. 3B). Both results suggest that the Tibeto-Burman-speaking migrants from the north and their admixture with local Austroasiatic speakers form the genetic profile of present-day Lahu and neighboring Tibeto-Burman speakers. We also observe that such a genetic profile had occurred in the Iron Age Thailand individuals ∼1,700 BP, with their genetic homogeneity with present-day Tibeto-Burman speakers in Mainland Southeast Asia confirmed by *qpWave* (Fig. 4A).

## DISCUSSION

In this study, we provide a comprehensive and detailed landscape for the genomic history of East Asians, especially Southern Chinese. We retrieve the deep origin and structure for the main ancestral groups in East Asia (Fig. 5A) and we document human migrations and admixtures that form the genomic and linguistic scenario in present-day Southern China (Fig. 5B). We predict that future ancient genomes from the interior of Southern China will further improve and examine the demographic framework of Southern East Asians established in our study.

## METHODS

### Sampling and genotyping

We collected blood and saliva samples from 211 unrelated individuals affiliated to Miao, Zhuang, and Han ethnicities from 30 subgroups in Guangxi and Yunnan of Southern China. Further linguistic and geographic information of these subgroups was described in Extended Data Table 1. The study was approved by Ethical Committee of Youjiang Medical University for Nationalities and all the processes involved were consistent with the corresponding ethical principles. All the participants read and signed the informed content. Then, we achieved the genotyped data of these samples using the Affymetrix WeGene V1 Array, which includes 492,683 genome-wide SNPs and is referred to as ‘500K dataset’ elsewhere in this paper. Other experimental and bioinformatic procedures for genotyping were consistent with the protocol documented in previous studies^38,39^.

### Dataset arrangement

We merged our 500K dataset with published present-day and ancient genomic data^6-9,11,12,17,18,24,32,33,39-56^, resulting in two types of panel: (1) merged panel of 500K dataset and 1240K-capture dataset (1,233,013 SNPs, including all the ancient samples and shotgun-sequenced modern samples) with 372,929 SNPs, which is for the purpose of maximizing the number of informative SNPs; (2) merged panel of the panel above and 600K Human Origin Array dataset (597,573 SNPs, including other modern samples) with 110,931 SNPs, which is for the purpose of maximizing the number and size of populations. For Rarecoal analysis, we used whole genome sequences from Simons Genome Diversity Project (SGDP)^47^.

### Principal component analysis (PCA)

We performed PCA by *smartpca* program of EIGENSOFT^13^ with parameters lsqproject: YES, shrinkmode: YES, numoutlieriter: 0, killr2: YES, r2thresh: 0.4, r2genlim: 0.1. We only used modern samples to construct PCs with ancient samples projected.

### ADMIXTURE analysis

We first used PLINK^57^ to prune the linkage disequilibrium by parameters --indep-pairwise 200 20 0.4. Then, we ran ADMIXTURE^14^ with default parameters from K = 2 to 20. We reported the result when K = 10 as it reaches the lowest cross error (Extended Data Fig. 2).

### *f*-statistics

We used ADMIXTOOLS^24^ to compute *f*_*3*_-statistics and D-statistics (Supplementary Information Table 2) with the estimation of standard error by jackknife. We used Mbuti as outgroup for Eurasian populations in outgroup-*f*_*3*_.

### Admixture graph modelling by *qpGraph*

We used *qpGraph* program of ADMIXTOOLS^24^ to reconstruct the phylogeny with admixture by default parameters. We exhausted different feasible graph models and select the optimal model based on maximum |Z|-score and likelihood.

### Admixture coefficient modelling by *qpAdm*

We used *qpAdm*^25^ to compute the ancestral coefficient based on f-statistics to different outgroups. We chose the optimal model for a given target population based on the following criteria, sorted by priority. (1) The model is feasible if and only if all the ancestral coefficients fall within the range [0, 1]. (2) The full model is chosen if both full and nested models are feasible. (3) If the full model is infeasible and more than one nested models are feasible, then the nested model with the highest *p*-value is chosen. We applied ‘proximal model’ and ‘distal model’^50^ to model the ancestry contribution in different time period.

#### Proximal model

We used Mongolia_N_East, Mekong_N (pooled population of Vietnam_N, Laos_LN_BA.SG, and Laos_BA.WGC), and Fujian_LN as proxies for Northern East Asian, Inland Southern East Asian, and Coastal Southern East Asian ancestries. The initial outgroups that we used are: South_Africa_2000BP.SG, Ust_Ishim.DG, Yana_UP.SG, Alaska_LP, Kolyma_M, Andaman_HG, Jomon_HG, Liangdao2_EN, Malaysia_LN.SG. We also used the ‘rotating’ strategy^41^ to further verify the nested models, in which we moved one of the proxies into the set of outgroups by turn. Since there is no high-coverage ancient sample that is sufficiently older than Mekong_N in Austroasiatic-related lineage, we expediently used Malaysia_LN.SG who closely related to Mekong_N as outgroup but we caution that it tends to underestimate *p*-values. Therefore, we also calculated relative likelihood ratios to test if a full model is better than its nested models and we find the ratios are usually higher than 100. Original results of proximal model are presented in Supplementary Information Table 1.

#### Distal model

We used Mongolia_N_East and Andaman_HG as proxies for East Asian and Andamanese-related ancestries. We used the following outgroups in distal model: South_Africa_2000BP.SG, Ust_Ishim.DG, Georgia_Kotias.SG, Loschbour.DG, Yana_UP.SG, Botai_EN, Russia_BA_Okunevo.SG, Russia_EHG_Karelia, Tianyuan, Papuan.DG, Mala.DG, Australian.DG, Hoabinhian.

### Genetic homogeneity testing by *qpWave*

We used *qpWave*^58^ to formally test if pairwise populations are homogeneous in relation to a series of outgroups. We used following outgroups for Southern East Asian populations: South_Africa_2000BP.SG, Ust_Ishim.DG, Loschbour.DG, Yana_UP.SG, Alaska_LP, Kolyma_M, Andaman_HG, Liangdao2_EN, Jomon_HG, Malaysia_LN.SG, Nepal_LN_BA_IA, DevilsCave_N, Shamanka_EN. We used following outgroups for Northern East Asian populations: South_Africa_2000BP.SG, Ust_Ishim.DG, Loschbour.DG, Yana_UP.SG, Alaska_LP, Kolyma_M, Andaman_HG, Liangdao2_EN, Jomon_HG, Malaysia_LN.SG, Nepal_LN_BA_IA, DevilsCave_N, Shamanka_EN.

### Demographic modelling implemented by Rarecoal

We used Rarecoal program^31,54^ to obtain a SFS-based phylogeny with time estimates using default parameters. We used mutation rate in every generation^59^ of 1.25 × 10^−8^ and 29 years per generation^60^ to scale the time.

### Admixture time estimation by ALDER

We used linkage disequilibrium-based ALDER^61^ to estimate admixture time of Hmong-Mien Cline using default parameters and checkmap: YES, mindis: 0.005, binsize: 0.0001.

### Identity-by-descent (IBD) analysis

We first used SHAPEIT^62^ to phase the modern individuals in our dataset. Then we used Refine IBD software^16^ to obtain pairwise sharing of IBD segments among individuals. We normalized the results in population level by dividing it by the product of the sample size of pairwise populations.

### Correlation between N_E_ and F_ST_ to Ust’-Ishim

We used the formula in Palamara et al.^63^ to estimate *N*_*E*_ from shared IBD within a population. We computed F_ST_ by *smartpca*^13^ with default parameters and fstonly: YES.

### LGM coastline in East Asia

The coastline during LGM period in East Asia shown in Fig. 5A is adopted from Ray et al..^64^

## Abbreviations

We used the following abbreviations throughout our article
LP: Late Pleistocene;
M: Mesolithic;
N: Neolithic;
EN: Early Neolithic;
MN: Middle Neolithic;
LN: Late Neolithic;
BA: Bronze Age;
IA: Iron Age;
o: outlier;
HG: hunter-gatherer;
MSEA: Mainland Southeast Asia;
ISEA: Island Southeast Asia;
AN: Austronesian;
AA: Austroasiatic;
HM: Hmong-Mien;
KD: Kra-Dai;
HO: Human Origin Array.
Particularly, Mongolia_N refers to Mongolia_N_East unless otherwise specified.

---

**Extended Data Figure 1.**
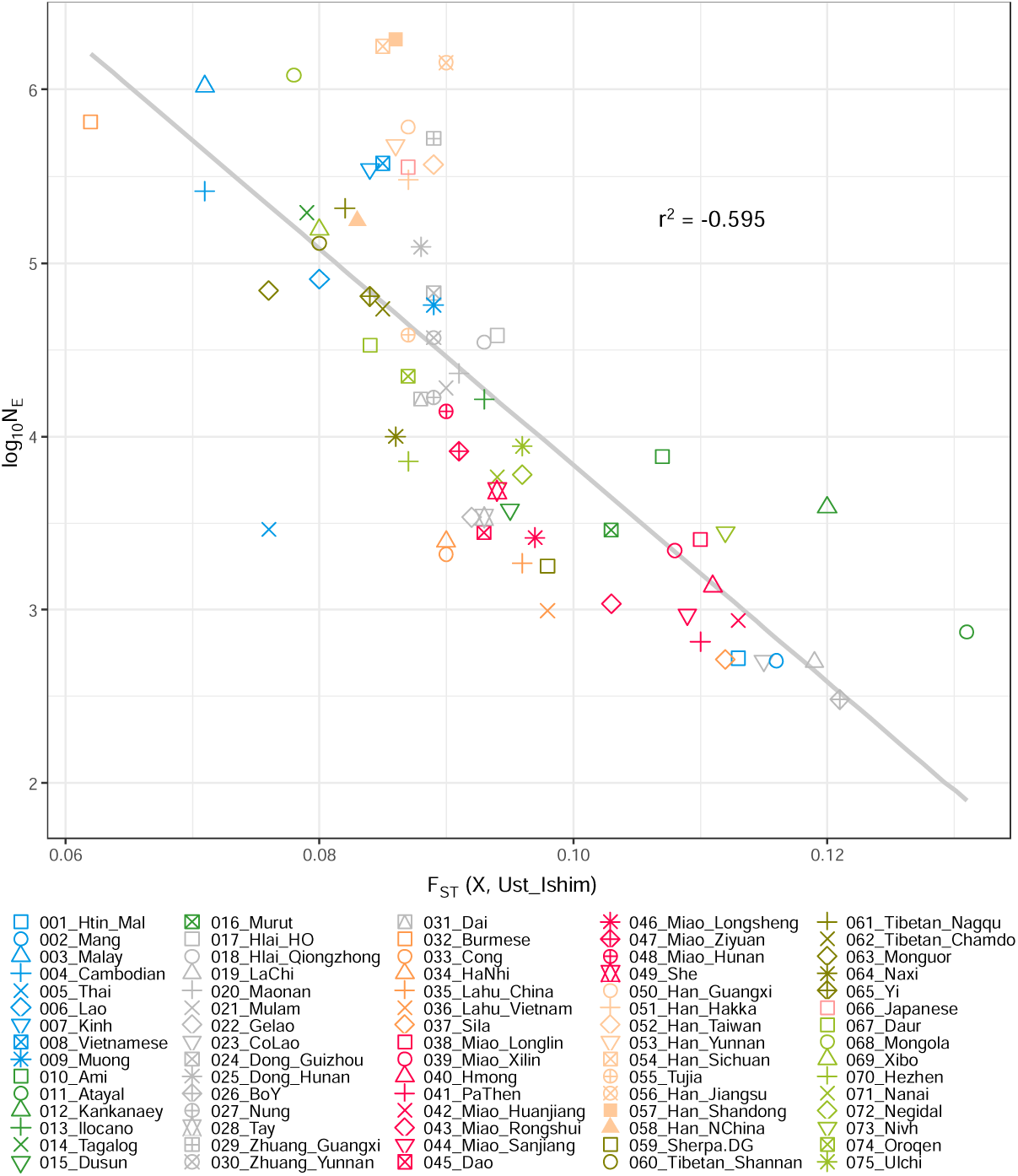
Correlation between effective population size (N_E_) and genetic drift away from common Eurasian ancestor. Given ∼45,000 years old Ust’-Ishim is genomically equally related to most of the Eurasians, we used F_ST_ away from him [F_ST_(X, Ust’-Ishim)] to represent the genetic drift from the common Eurasian ancestor to modern East Asian populations. Negative correlation between logarithm of N_E_ and F_ST_(X, Ust’-Ishim) suggests that recent genetic drift due to a small population size comprise a large proportion of the total genetic drift from common Eurasian ancestor in many modern East Asian populations.

**Extended Data Figure 2.**
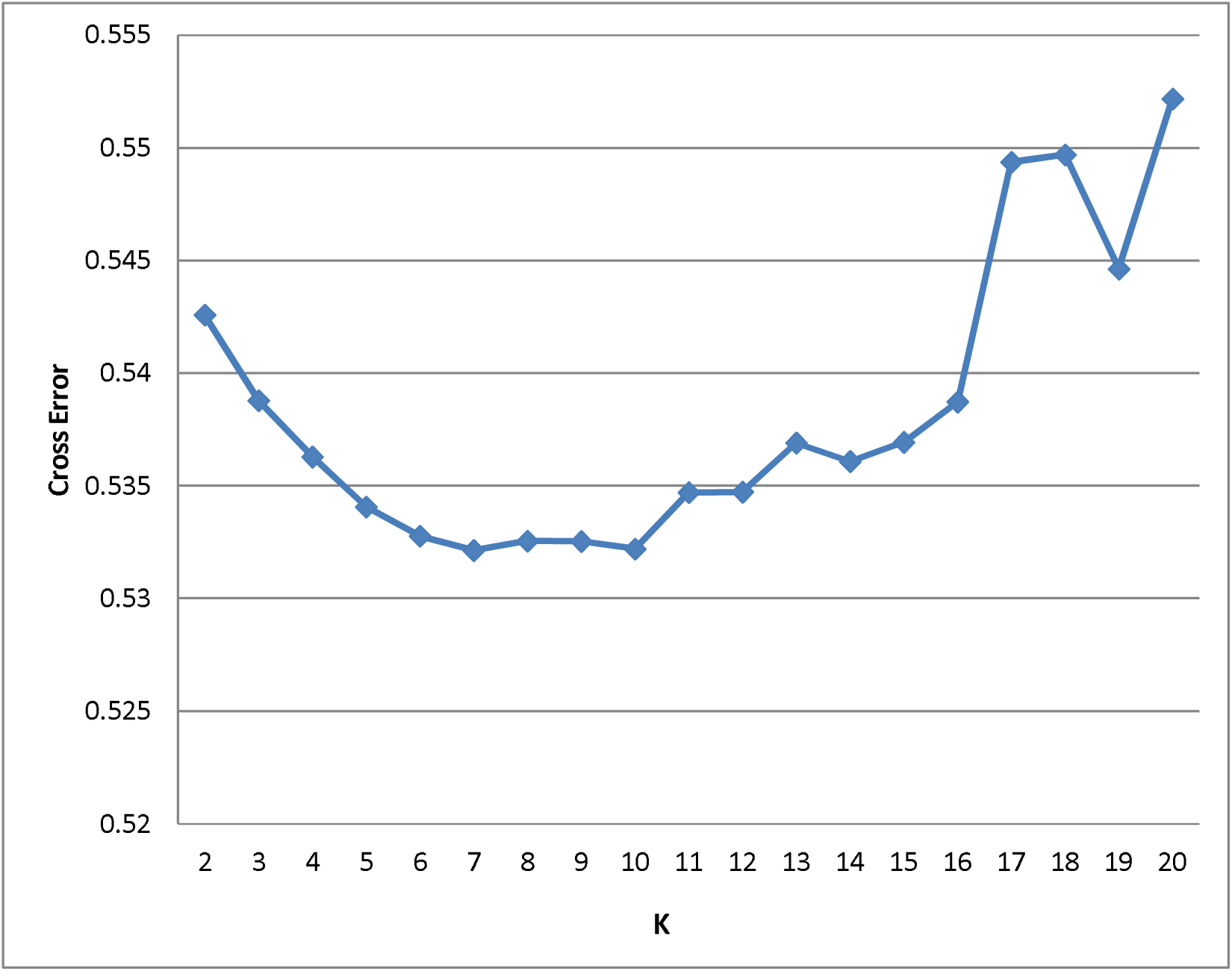
Cross error for ADMIXTURE analysis when K = 2 to 20.

**Extended Data Table 1.** Sample information for newly reported individuals in this study.

| Population | Language Affiliation | Locality | Region | Latitude | Longitude | Grouped label | Sample size |
| --- | --- | --- | --- | --- | --- | --- | --- |
| Miao_Longlin | HM, Hmongic, Hmong | Longlin, Baise, Guangxi | China (Southwest) | 24.7714 | 105.3456 | Miao_Longlin | 10 |
| Miao_Xilin | HM, Hmongic, Hmong | Xilin, Baise, Guangxi | China (Southwest) | 24.4924 | 105.0929 | Miao_Xilin | 10 |
| Miao_Huanjiang | HM, Hmongic, Hmu, Southern Dialect | Huanjiang, Hechi, Guangxi | China (Southwest) | 24.8363 | 108.2538 | Miao_Huanjiang | 8 |
| Miao_Rongshui | HM, Hmongic, Hmu, Southern Dialect | Rongshui, Liuzhou, Guangxi | China (Southwest) | 25.0748 | 109.2537 | Miao_Rongshui | 10 |
| Miao_Sanjiang | HM, Hmongic, Hmu, Southern Dialect | Sanjiang, Liuzhou, Guangxi | China (Southwest) | 25.7818 | 109.6064 | Miao_Sanjiang | 9 |
| Miao_Ziyuan | HM, Hmongic, Hmu, Eastern Dialect | Ziyuan, Guilin, Guangxi | China (Southwest) | 26.0365 | 110.6397 | Miao_Ziyuan | 10 |
| Miao_Longsheng | HM, Hmongic, Hmu, Eastern Dialect | Longsheng, Guilin, Guangxi | China (Southwest) | 25.7992 | 110.0081 | Miao_Longsheng | 10 |
| Zhuang_Qiubei | KD, Tai, Northern Tai | Qiubei, Wenshan, Yunnan | China (Southwest) | 24.0428 | 104.1887 | Zhuang_Yunnan | 9 |
| Zhuang_Guangnan | KD, Tai, Central Tai | Guangnan, Wenshan, Yunnan | China (Southwest) | 24.0458 | 105.0535 | Zhuang_Yunnan | 5 |
| Zhuang_Wenshan | KD, Tai, Central Tai | Wenshan, Wenshan, Yunnan | China (Southwest) | 23.3864 | 104.2318 | Zhuang_Yunnan | 10 |
| Zhuang_Tianlin | KD, Tai, Northern Tai | Tianlin, Baise, Guangxi | China (Southwest) | 24.2946 | 106.2306 | Zhuang_Guangxi | 10 |
| Zhuang_Tianyang | KD, Tai, Northern Tai | Tianyang, Baise, Guangxi | China (Southwest) | 23.7377 | 106.9160 | Zhuang_Guangxi | 6 |
| Zhuang_Jingxi | KD, Tai, Central Tai | Jingxi, Baise, Guangxi | China (Southwest) | 23.1355 | 106.4171 | Zhuang_Guangxi | 9 |
| Zhuang_Chongzuo | KD, Tai, Central Tai | Jiangzhou, Chongzuo, Guangxi | China (Southwest) | 22.4071 | 107.3546 | Zhuang_Guangxi | 10 |
| Zhuang_Fusui | KD, Tai, Central Tai | Fusui, Chongzuo, Guangxi | China (Southwest) | 22.6350 | 107.9041 | Zhuang_Guangxi | 6 |
| Zhuang_Wuming | KD, Tai, Central Tai | Wuming, Nanning, Guangxi | China (Southwest) | 23.1560 | 108.2841 | Zhuang_Guangxi | 1 |
| Zhuang_Hechi | KD, Tai, Northern Tai | Jinchengjiang, Hechi, Guangxi | China (Southwest) | 24.6944 | 108.0846 | Zhuang_Guangxi | 9 |
| Zhuang_Guigang | KD, Tai, Northern Tai | Gangnan, Guigang, Guangxi | China (Southwest) | 23.1107 | 109.5978 | Zhuang_Guangxi | 10 |
| Zhuang_Laibin | KD, Tai, Northern Tai | Xingbin, Laibin, Guangxi | China (Southwest) | 23.7402 | 109.1962 | Zhuang_Guangxi | 6 |
| Han_Qiubei | ST, Sinitic, SW Mandarin, Diannan | Qiubei, Wenshan, Yunnan | China (Southwest) | 24.0428 | 104.1887 | Han_Yunnan | 6 |
| Han_Guangnan | ST, Sinitic, SW Mandarin, Diannan | Guangnan, Wenshan, Yunnan | China (Southwest) | 24.0458 | 105.0535 | Han_Yunnan | 3 |
| Han_Wenshan | ST, Sinitic, SW Mandarin, Diannan | Wenshan, Wenshan, Yunnan | China (Southwest) | 23.3864 | 104.2318 | Han_Yunnan | 7 |
| Han_Tianlin | ST, Sinitic, Pinghua | Tianlin, Baise, Guangxi | China (Southwest) | 24.2946 | 106.2306 | Han_Guangxi | 5 |
| Han_Tianyang | ST, Sinitic, Pinghua | Tianyang, Baise, Guangxi | China (Southwest) | 23.7377 | 106.9160 | Han_Guangxi | 5 |
| Han_Jingxi | ST, Sinitic, SW Mandarin, Guliu | Jingxi, Baise, Guangxi | China (Southwest) | 23.1355 | 106.4171 | Han_Guangxi | 5 |
| Han_Chongzuo | ST, Sinitic, Pinghua | Jiangzhou, Chongzuo, Guangxi | China (Southwest) | 22.4071 | 107.3546 | Han_Guangxi | 5 |
| Han_Fusui | ST, Sinitic, Pinghua | Fusui, Chongzuo, Guangxi | China (Southwest) | 22.6350 | 107.9041 | Han_Guangxi | 1 |
| Han_Hechi | ST, Sinitic, SW Mandarin, Guliu | Jinchengjiang, Hechi, Guangxi | China (Southwest) | 24.6944 | 108.0846 | Han_Guangxi | 6 |
| Han_Guigang | ST, Sinitic, Pinghua | Gangnan, Guigang, Guangxi | China (Southwest) | 23.1107 | 109.5978 | Han_Guangxi | 6 |
| Han_Laibin | ST, Sinitic, SW Mandarin, Guliu | Xingbin, Laibin, Guangxi | China (Southwest) | 23.7402 | 109.1962 | Han_Guangxi | 4 |

**Extended Data Table 2.** Correlation between the proportion of ancestries and corresponding language families. We used point-biserial correlation to quantify if an individual affiliated to a certain language family tends to have more proportion of the ancestry corresponding to this language family. We further used the *p*-value of student’s t-test to quantify if the correlation is significant. r_pb_, point-biserial correlation coefficient.

| **Language Families** | **$r_{pb}$** | **$p$-value** |
| --- | --- | --- |
| Hmong-Mien | 0.762 | $2.13 \times 10^{-31}$ |
| Kra-Dai | 0.644 | $9.77 \times 10^{-189}$ |
| Austroasiatic | 0.536 | $6.73 \times 10^{-12}$ |
| Austronesian | 0.921 | $7.23 \times 10^{-32}$ |
| Sino-Tibetan | 0.630 | $4.81 \times 10^{-83}$ |
| Tungusic/ Amuric | 0.806 | $7.34 \times 10^{-31}$ |

**Extended Data Table 3.** Admixture-*f*_*3*_ results. (**A)** Tibeto-Burman populations. (**B**) Southeast Han Chinese. (C) Kra-Dai and Vietic populations. We report the five lowest *f*_*3*_ results for each of the populations. std.err, standard error.

**A**
| Source_1 | Source_2 | Target | $f_3$ | std.err | Z | SNPs |
| --- | --- | --- | --- | --- | --- | --- |
| Mekong_N | Yumin_EN | Naxi | -0.001575 | 0.002227 | -0.708 | 69321 |
| Mlabri | Tibetan_Chamdo | Naxi | -0.001207 | 0.000555 | -2.177 | 105215 |
| Mekong_N | Sherpa.DG | Naxi | -0.001042 | 0.001291 | -0.807 | 83146 |
| Mekong_N | Tibetan_Chamdo | Naxi | -0.001036 | 0.00068 | -1.523 | 87697 |
| Mekong_N | Tibetan_Nagqu | Naxi | -0.000908 | 0.000777 | -1.17 | 87061 |
| Mlabri | Tibetan_Chamdo | Yi | -0.00375 | 0.000573 | -6.544 | 105393 |
| Malaysia_LN.SG | Yumin_EN | Yi | -0.003742 | 0.002493 | -1.501 | 67896 |
| Mekong_N | Yumin_EN | Yi | -0.003681 | 0.002139 | -1.721 | 70135 |
| Ami | Tibetan_Chamdo | Yi | -0.00366 | 0.000363 | -10.085 | 106865 |
| Malaysia_LN.SG | Tibetan_Chamdo | Yi | -0.003537 | 0.000795 | -4.451 | 84607 |
| Mekong_N | Yumin_EN | Sila | 0.029056 | 0.002478 | 11.724 | 67679 |
| Mekong_N | Qijia_LN | Sila | 0.030261 | 0.001622 | 18.661 | 81896 |
| Malaysia_LN.SG | Qijia_LN | Sila | 0.03054 | 0.001782 | 17.139 | 78608 |
| Mlabri | Tibetan_Chamdo | Sila | 0.03056 | 0.000988 | 30.924 | 104727 |
| Mekong_N | Tibetan_Chamdo | Sila | 0.030608 | 0.001149 | 26.649 | 87209 |
| Mekong_N | Yumin_EN | HaNhi | 0.00217 | 0.002201 | 0.986 | 69776 |
| Mlabri | Tibetan_Chamdo | HaNhi | 0.003446 | 0.000648 | 5.317 | 105256 |
| Mekong_N | Tibetan_Nagqu | HaNhi | 0.003451 | 0.000859 | 4.018 | 87152 |
| Mlabri | Lokomotiv_EN | HaNhi | 0.003579 | 0.001226 | 2.919 | 89625 |
| Mekong_N | Qijia_LN | HaNhi | 0.00363 | 0.001297 | 2.799 | 83933 |
| Mekong_N | Yumin_EN | Cong | 0.003042 | 0.002203 | 1.381 | 69905 |
| Malaysia_LN.SG | Qijia_LN | Cong | 0.00389 | 0.001518 | 2.563 | 80814 |
| Mekong_N | Sherpa.DG | Cong | 0.004221 | 0.00135 | 3.127 | 83654 |
| Mekong_N | Qijia_LN | Cong | 0.004291 | 0.001311 | 3.274 | 84076 |
| Malaysia_LN.SG | Yumin_EN | Cong | 0.004553 | 0.002551 | 1.785 | 67698 |
| Mekong_N | Yumin_EN | Lahu_China | 0.009174 | 0.002329 | 3.94 | 67477 |
| Malaysia_LN.SG | Yumin_EN | Lahu_China | 0.010163 | 0.002762 | 3.68 | 65140 |
| Malaysia_LN.SG | Qijia_LN | Lahu_China | 0.010199 | 0.001622 | 6.286 | 78604 |
| Malaysia_LN.SG | Tibetan_Chamdo | Lahu_China | 0.010683 | 0.001132 | 9.437 | 84250 |
| Mekong_N | Sherpa.DG | Lahu_China | 0.010862 | 0.001495 | 7.267 | 81702 |
| Mekong_N | Yumin_EN | Lahu_Vietnam | 0.008825 | 0.00227 | 3.889 | 68785 |
| Malaysia_LN.SG | Yumin_EN | Lahu_Vietnam | 0.010852 | 0.002663 | 4.074 | 66561 |
| Mekong_N | Qijia_LN | Lahu_Vietnam | 0.012234 | 0.001375 | 8.895 | 82996 |
| Mlabri | Yumin_EN | Lahu_Vietnam | 0.012279 | 0.001718 | 7.146 | 84001 |
| Malaysia_LN.SG | Qijia_LN | Lahu_Vietnam | 0.012334 | 0.001543 | 7.994 | 79719 |

B
| Source_1 | Source_2 | Target | f_3 | std.err | Z | SNPs |
| --- | --- | --- | --- | --- | --- | --- |
| Atayal | Qijia_LN | Han_Fujian | -0.005661 | 0.000819 | -6.912 | 100135 |
| Kankanaey | Qijia_LN | Han_Fujian | -0.004939 | 0.000759 | -6.506 | 100676 |
| Atayal | Yumin_EN | Han_Fujian | -0.004855 | 0.00132 | -3.68 | 83301 |
| Hlai_Qiongzong | Qijia_LN | Han_Fujian | -0.004405 | 0.000645 | -6.828 | 105279 |
| Kankanaey | Yangshao_MN | Han_Fujian | -0.004354 | 0.000834 | -5.222 | 100270 |
| Atayal | Yumin_EN | Han_Guangdong | -0.005255 | 0.001198 | -4.385 | 84603 |
| Hlai_Qiongzong | Yumin_EN | Han_Guangdong | -0.004742 | 0.000835 | -5.677 | 88521 |
| Atayal | Qijia_LN | Han_Guangdong | -0.004456 | 0.000746 | -5.97 | 101411 |
| Ami | Yumin_EN | Han_Guangdong | -0.004278 | 0.001073 | -3.989 | 85412 |
| Maonan | Yumin_EN | Han_Guangdong | -0.004173 | 0.000876 | -4.762 | 87806 |
| Atayal | Qijia_LN | Han_Taiwan | -0.004181 | 0.000711 | -5.882 | 101169 |
| Atayal | Yumin_EN | Han_Taiwan | -0.004013 | 0.001229 | -3.266 | 84541 |
| Hlai_Qiongzong | Yumin_EN | Han_Taiwan | -0.003404 | 0.000833 | -4.084 | 88057 |
| Kankanaey | Qijia_LN | Han_Taiwan | -0.00315 | 0.000651 | -4.836 | 101443 |
| Hlai_Qiongzong | Qijia_LN | Han_Taiwan | -0.003081 | 0.000511 | -6.034 | 104904 |
| Qijia_LN | Atayal | Han_Hakka | -0.005343 | 0.000696 | -7.673 | 101131 |
| Qijia_LN | Kankanaey | Han_Hakka | -0.004676 | 0.000635 | -7.367 | 101487 |
| Sherpa.DG | Kankanaey | Han_Hakka | -0.004664 | 0.000734 | -6.351 | 100941 |
| Tibetan_Chamdo | Ami | Han_Hakka | -0.004617 | 0.000436 | -10.584 | 105655 |
| Sherpa.DG | Ami | Han_Hakka | -0.004606 | 0.00068 | -6.774 | 101421 |
| Mongolia_N_North | Hlai_HO | Han_Guangxi | -0.004074 | 0.000537 | -7.583 | 103164 |
| Qijia_LN | Atayal | Han_Guangxi | -0.004072 | 0.00054 | -7.54 | 107035 |
| Yumin_EN | Atayal | Han_Guangxi | -0.003639 | 0.000981 | -3.71 | 89909 |
| Qijia_LN | Hlai_HO | Han_Guangxi | -0.003628 | 0.000494 | -7.344 | 106925 |
| Qijia_LN | Hlai_Qiongzong | Han_Guangxi | -0.003512 | 0.00029 | -12.119 | 107881 |
| Atayal | Qijia_LN | Han_Zhejiang | -0.005071 | 0.000843 | -6.015 | 100136 |
| Kankanaey | Qijia_LN | Han_Zhejiang | -0.004388 | 0.0008 | -5.486 | 100775 |
| Ami | Qijia_LN | Han_Zhejiang | -0.004223 | 0.000729 | -5.797 | 101373 |
| Hlai_HO | Qijia_LN | Han_Zhejiang | -0.004132 | 0.000812 | -5.089 | 99336 |
| Hlai_HO | Boisman_MN | Han_Zhejiang | -0.003947 | 0.000837 | -4.717 | 93260 |
| Atayal | Qijia_LN | CHS.SG | -0.0021 | 0.000535 | -3.927 | 107824 |
| Kankanaey | Qijia_LN | CHS.SG | -0.0013 | 0.000469 | -2.774 | 107480 |
| Ami | Qijia_LN | CHS.SG | -0.000939 | 0.000421 | -2.23 | 107882 |
| Hlai_Qiongzong | Qijia_LN | CHS.SG | -0.00088 | 0.0003 | -2.936 | 108513 |
| Hlai_HO | Qijia_LN | CHS.SG | -0.000867 | 0.000481 | -1.802 | 107768 |

C
| Source_1 | Source_2 | Target | f_3 | std.err | Z | SNPs |
| --- | --- | --- | --- | --- | --- | --- |
| Mlabri | Fujian_LN | Hlai_Qiongzong | 0.001787 | 0.0011920 | 1.499 | 69995 |
| Mekong_N | Ami | Hlai_Qiongzong | 0.002454 | 0.0007040 | 3.485 | 87834 |
| Mekong_N | Atayal | Hlai_Qiongzong | 0.003009 | 0.0008930 | 3.370 | 87595 |
| Malaysia_LN.SG | Ami | Hlai_Qiongzong | 0.003022 | 0.0008760 | 3.449 | 84545 |
| Mekong_N | Kankanaey | Hlai_Qiongzong | 0.003176 | 0.0008100 | 3.920 | 87457 |
| Mlabri | Fujian_LN | Zhuang_Guangxi | -0.002579 | 0.0011180 | -2.306 | 72220 |
| Hlai_HO | Mongolia_N_North | Zhuang_Guangxi | -0.002157 | 0.0005480 | -3.934 | 104259 |
| Mekong_N | Ami | Zhuang_Guangxi | -0.002070 | 0.0006260 | -3.309 | 90401 |
| Hlai_HO | Kolyma_M | Zhuang_Guangxi | -0.001675 | 0.0008670 | -1.932 | 108675 |
| Mekong_N | Atayal | Zhuang_Guangxi | -0.001607 | 0.0008380 | -1.917 | 90347 |
| Mlabri | Fujian_LN | Zhuang_Yunnan | -0.002457 | 0.001131 | -2.172 | 70278 |
| Mlabri | Qihe_EN | Zhuang_Yunnan | -0.00186 | 0.001865 | -0.998 | 38807 |
| Mlabri | Kankanaey | Zhuang_Yunnan | -0.001246 | 0.000609 | -2.045 | 105224 |
| Mekong_N | Atayal | Zhuang_Yunnan | -0.001242 | 0.00091 | -1.365 | 88055 |
| Mekong_N | Ami | Zhuang_Yunnan | -0.001209 | 0.000707 | -1.709 | 88247 |
| Mlabri | Fujian_LN | CDX.SG | -0.00053 | 0.001087 | -0.487 | 72015 |
| Mlabri | Qihe_EN | CDX.SG | -0.000412 | 0.001793 | -0.23 | 39820 |
| Mlabri | Liangdao1_EN | CDX.SG | 0.000201 | 0.001681 | 0.119 | 56649 |
| Mekong_N | Ami | CDX.SG | 0.00031 | 0.000664 | 0.466 | 90185 |
| Mekong_N | Hezhen | CDX.SG | 0.000581 | 0.000696 | 0.835 | 90781 |
| Hlai_HO | Kolyma_M | Mulam | -0.000988 | 0.000961 | -1.028 | 104445 |
| Hlai_HO | Mongolia_N_North | Mulam | -0.000853 | 0.000629 | -1.356 | 100764 |
| Fujian_LN | Mlabri | Mulam | -0.000814 | 0.001178 | -0.692 | 69313 |
| Hlai_HO | Qijia_LN | Mulam | -0.000519 | 0.000571 | -0.909 | 104153 |
| Hlai_Qiongzong | Qijia_LN | Mulam | -0.000489 | 0.000364 | -1.343 | 106533 |
| Mlabri | Fujian_LN | Dai | -0.001628 | 0.001278 | -1.273 | 67817 |
| Mekong_N | Atayal | Dai | -0.001387 | 0.000958 | -1.447 | 85437 |
| Mekong_N | Ami | Dai | -0.001195 | 0.000813 | -1.47 | 86033 |
| Mlabri | Liangdao1_EN | Dai | -0.001099 | 0.001813 | -0.606 | 53082 |
| Mekong_N | Kankanaey | Dai | -0.000769 | 0.000895 | -0.858 | 85539 |
| Qihe_EN | Mlabri | Maonan | -0.000708 | 0.001897 | -0.373 | 38205 |
| Fujian_LN | Mlabri | Maonan | -0.000669 | 0.001211 | -0.553 | 69171 |
| Hlai_HO | Kolyma_M | Maonan | -0.000049 | 0.000991 | -0.05 | 104275 |
| Qihe_EN | Miao_Longlin | Maonan | 0.000224 | 0.0012 | 0.187 | 38806 |
| Ami | Mekong_N | Maonan | 0.000309 | 0.000745 | 0.414 | 87016 |
| Miao_Longlin | Qihe_EN | CoLao | 0.026857 | 0.001809 | 14.85 | 37582 |
| Miao_Longlin | Liangdao1_EN | CoLao | 0.02686 | 0.001497 | 17.938 | 53446 |
| Miao_Xilin | Malaysia_LN.SG | CoLao | 0.027253 | 0.001376 | 19.812 | 81187 |
| Miao_Longlin | Mekong_N | CoLao | 0.02745 | 0.001196 | 22.945 | 84419 |
| Yumin_EN | Mekong_N | CoLao | 0.027586 | 0.002572 | 10.727 | 66989 |
| Mlabri | Qihe_EN | Gelao | 0.000421 | 0.001948 | 0.216 | 37203 |
| Mlabri | Fujian_LN | Gelao | 0.00056 | 0.001252 | 0.447 | 67550 |
| Mlabri | Longshan_LN | Gelao | 0.001273 | 0.000756 | 1.683 | 102065 |
| Mlabri | Yangshao_MN | Gelao | 0.001494 | 0.001025 | 1.457 | 100619 |
| Mlabri | Liangdao1_EN | Gelao | 0.001592 | 0.001851 | 0.86 | 52851 |
| Mlabri | Fujian_LN | LaChi | 0.033654 | 0.00162 | 20.769 | 66235 |
| Malaysia_LN.SG | Qijia_LN | LaChi | 0.034132 | 0.001792 | 19.042 | 78138 |
| Mekong_N | Ami | LaChi | 0.034462 | 0.001275 | 27.033 | 84541 |
| Malaysia_LN.SG | Yumin_EN | LaChi | 0.034524 | 0.002895 | 11.925 | 64779 |
| Mlabri | Qihe_EN | LaChi | 0.034846 | 0.002477 | 14.07 | 36410 |
| Malaysia_LN.SG | Yumin_EN | Kinh | -0.005497 | 0.002521 | -2.18 | 66590 |
| Mekong_N | Yumin_EN | Kinh | -0.004957 | 0.002151 | -2.304 | 68862 |
| Mlabri | Fujian_LN | Kinh | -0.00494 | 0.001269 | -3.892 | 67451 |
| Malaysia_LN.SG | Qijia_LN | Kinh | -0.004437 | 0.001486 | -2.985 | 80021 |
| Mlabri | Yangshao_MN | Kinh | -0.004388 | 0.001001 | -4.382 | 100543 |
| Mlabri | Qihe_EN | Muong | -0.003278 | 0.001942 | -1.688 | 37341 |
| Mlabri | Fujian_LN | Muong | -0.002737 | 0.001205 | -2.273 | 67833 |
| Mekong_N | Ami | Muong | -0.002661 | 0.000843 | -3.158 | 85867 |
| Mekong_N | Kankanaey | Muong | -0.00232 | 0.000908 | -2.553 | 85724 |
| Mekong_N | Atayal | Muong | -0.002275 | 0.000966 | -2.354 | 85258 |
| Mekong_N | Hezhen | KHV.SG | -0.001114 | 0.000653 | -1.705 | 91653 |
| Mekong_N | Yumin_EN | KHV.SG | -0.001059 | 0.002048 | -0.517 | 75980 |
| Malaysia_LN.SG | Qijia_LN | KHV.SG | -0.001 | 0.001286 | -0.778 | 87386 |
| Mlabri | Qihe_EN | KHV.SG | -0.000881 | 0.001796 | -0.491 | 40278 |
| Mlabri | Fujian_LN | KHV.SG | -0.000864 | 0.001083 | -0.798 | 72809 |
| Mekong_N | Ami | Nung | -0.001796 | 0.000768 | -2.339 | 85969 |
| Mekong_N | Atayal | Nung | -0.00178 | 0.000917 | -1.941 | 85459 |
| Mlabri | Fujian_LN | Nung | -0.001548 | 0.00118 | -1.311 | 68167 |
| Mekong_N | Hezhen | Nung | -0.001542 | 0.000792 | -1.946 | 87392 |
| Mlabri | Qihe_EN | Nung | -0.001188 | 0.001907 | -0.623 | 37606 |
| Mlabri | Qihe_EN | Lao | -0.006126 | 0.001913 | -3.203 | 37733 |
| Mlabri | Fujian_LN | Lao | -0.006086 | 0.00118 | -5.158 | 68494 |
| Mekong_N | Ami | Lao | -0.005638 | 0.000799 | -7.057 | 86114 |
| Mlabri | Liangdao1_EN | Lao | -0.005427 | 0.001802 | -3.012 | 53597 |
| Mekong_N | Hezhen | Lao | -0.005185 | 0.000809 | -6.41 | 87452 |

**Extended Data Table 4.** Admixture time estimates for Hmong-Mien Cline inferred by ALDER.

| **p-value** | **target** | **reference A** | **reference B** | **Z-score** | **admixture time estimate** |
| --- | --- | --- | --- | --- | --- |
| 1.5E-10 | Miao_Rongshui | Hmong_Core | CHB.SG | 6.40 | 24.43 ± 3.82 |
| 1.6E-07 | Miao_Sanjiang | Hmong_Core | Austronesian_Core | 5.25 | 25.80 ± 4.92 |
| 6.8E-09 | Miao_Rongshui | Hmong_Core | KHV.SG | 5.80 | 25.92 ± 4.47 |
| 1.8E-13 | Miao_Rongshui | Hmong_Core | CDX.SG | 7.36 | 26.76 ± 3.64 |
| 4.4E-10 | Miao_Sanjiang | Hmong_Core | Tibetan_Core | 6.24 | 26.86 ± 4.30 |
| 4.8E-07 | Dong_Hunan | Hmong_Core | Hlai_all | 5.03 | 26.97 ± 5.36 |
| 5.4E-09 | Miao_Rongshui | Hmong_Core | Amur_Core | 5.83 | 27.02 ± 4.63 |
| 2.9E-09 | Miao_Sanjiang | Hmong_Core | CHS.SG | 5.93 | 27.08 ± 4.56 |
| 2.9E-09 | Miao_Sanjiang | Hmong_Core | CHS.SG | 5.93 | 27.08 ± 4.56 |
| 2.8E-06 | Dong_Hunan | Hmong_Core | CHB.SG | 4.69 | 28.36 ± 6.05 |
| 1.0E-06 | Dong_Hunan | Hmong_Core | Tibetan_Core | 4.89 | 29.32 ± 6.00 |
| 4.9E-08 | Miao_Sanjiang | Hmong_Core | CDX.SG | 5.46 | 29.33 ± 5.38 |
| 1.8E-06 | Dong_Hunan | Hmong_Core | KHV.SG | 4.78 | 29.38 ± 6.15 |
| 1.3E-07 | Dong_Hunan | Hmong_Core | CHS.SG | 5.28 | 29.91 ± 5.67 |
| 1.3E-07 | Dong_Hunan | Hmong_Core | CHS.SG | 5.28 | 29.91 ± 5.67 |
| 7.5E-18 | Miao_Huanjiang | Hmong_Core | Hlai_all | 8.61 | 31.09 ± 3.61 |
| 2.3E-12 | Dong_Guizhou | Hmong_Core | KHV.SG | 7.01 | 31.97 ± 4.56 |
| 5.7E-19 | Miao_Huanjiang | Hmong_Core | KHV.SG | 8.90 | 32.09 ± 3.61 |
| 2.5E-18 | Miao_Huanjiang | Hmong_Core | CHB.SG | 8.73 | 33.91 ± 3.88 |
| 2.1E-07 | Dong_Guizhou | Hmong_Core | CDX.SG | 5.19 | 34.58 ± 6.67 |
| 1.6E-18 | Miao_Huanjiang | Hmong_Core | Zhuang_Guangxi | 8.79 | 35.01 ± 3.98 |
| 1.3E-09 | PaThen | Hmong_Core | Zhuang_Guangxi | 6.07 | 37.30 ± 6.14 |
| 1.9E-09 | Dong_Guizhou | Hmong_Core | CHB.SG | 6.01 | 38.26 ± 6.37 |
| 1.4E-15 | Dong_Guizhou | Hmong_Core | CHS.SG | 7.98 | 38.91 ± 4.87 |
| 1.4E-15 | Dong_Guizhou | Hmong_Core | CHS.SG | 7.98 | 38.91 ± 4.87 |
| 3.0E-10 | PaThen | Hmong_Core | CHS.SG | 6.30 | 39.73 ± 6.31 |
| 3.0E-10 | PaThen | Hmong_Core | CHS.SG | 6.30 | 39.73 ± 6.31 |
| 3.8E-09 | PaThen | Hmong_Core | CHB.SG | 5.89 | 40.25 ± 6.83 |
| 1.5E-09 | Dong_Guizhou | Hmong_Core | JPT.SG | 6.04 | 40.27 ± 6.67 |
| 7.2E-19 | Miao_Huanjiang | Hmong_Core | Tibetan_Core | 8.87 | 40.86 ± 4.61 |
| 1.2E-08 | Dong_Guizhou | Hmong_Core | Tibetan_Core | 5.70 | 41.91 ± 7.09 |
| 1.7E-05 | Miao_Longsheng | Hmong_Core | KHV.SG | 4.30 | 41.91 ± 9.74 |
| 4.1E-05 | Miao_Longsheng | Hmong_Core | CHS.SG | 4.10 | 45.62 ± 11.13 |
| 4.1E-05 | Miao_Longsheng | Hmong_Core | CHS.SG | 4.10 | 45.62 ± 11.13 |
| 4.6E-05 | Miao_Longsheng | Hmong_Core | CDX.SG | 4.08 | 46.40 ± 11.38 |

**Extended Data Table 5.**

| Pop | P-value | Ancestry Coefficient |  | Standard Error |  |
| --- | --- | --- | --- | --- | --- |
|  |  | Mongolia_N | Andaman_HG | Mongolia_N | Andaman_HG |
| Haminmangha_MN | 2.139E-01 | 0.852 | 0.148 | 0.065 | 0.065 |
| Jalainur_EN | 9.769E-01 | 0.992 | 0.008 | 0.072 | 0.072 |
| Xianbei_IA | 4.856E-02 | 0.956 | 0.044 | 0.092 | 0.092 |
| Heishui_Mohe_Medieval | 4.749E-01 | 0.707 | 0.293 | 0.073 | 0.073 |
| Yankovsky_IA | 2.113E-01 | 0.969 | 0.031 | 0.060 | 0.060 |
| Boisman_MN | 5.588E-01 | 0.936 | 0.064 | 0.040 | 0.040 |
| DevilsCave_N | 3.318E-01 | 0.883 | 0.117 | 0.042 | 0.042 |
| U_Xiajiadian_BA_o | 7.984E-02 | 1.046 | -0.046 | 0.065 | 0.065 |
| Yumin_EN | 4.060E-02 | 0.978 | 0.022 | 0.058 | 0.058 |
| Mongolia_N_North | 6.086E-02 | 1.025 | -0.025 | 0.043 | 0.043 |
| Hongshan_MN | 7.480E-01 | 0.869 | 0.131 | 0.044 | 0.044 |
| L_Xiajiadian_LN | 6.424E-01 | 0.837 | 0.163 | 0.045 | 0.045 |
| U_Xiajiadian_BA | 4.833E-01 | 0.900 | 0.100 | 0.057 | 0.057 |
| Jomon_HG | 9.088E-01 | 0.435 | 0.565 | 0.046 | 0.046 |
| Sakhalin_HG | 1.574E-01 | 0.552 | 0.448 | 0.116 | 0.116 |
| Shandong_EN | 3.289E-01 | 0.861 | 0.139 | 0.037 | 0.037 |
| Yangshao_MN | 9.339E-01 | 0.787 | 0.213 | 0.038 | 0.038 |
| Longshan_LN | 6.039E-01 | 0.826 | 0.174 | 0.033 | 0.033 |
| Miaozigou_MN | 8.369E-01 | 0.922 | 0.078 | 0.076 | 0.076 |
| Shimao_LN | 1.909E-01 | 0.853 | 0.147 | 0.047 | 0.047 |
| Henan_LBIA | 3.191E-01 | 0.797 | 0.203 | 0.034 | 0.034 |
| Omnogovi_WestHan_IA | 5.448E-01 | 0.786 | 0.214 | 0.042 | 0.042 |
| Qijia_LN | 4.798E-01 | 0.780 | 0.220 | 0.038 | 0.038 |
| Dacaozi_IA | 2.658E-01 | 0.820 | 0.180 | 0.044 | 0.044 |
| Nepal_LN_BA_IA | 9.297E-02 | 0.702 | 0.298 | 0.038 | 0.038 |
| Laos_N | 6.922E-01 | 0.442 | 0.558 | 0.050 | 0.050 |
| Malaysia_LN.SG | 3.746E-01 | 0.483 | 0.517 | 0.050 | 0.050 |
| Vietnam_N | 8.132E-01 | 0.426 | 0.574 | 0.105 | 0.105 |
| Vietnam_LN_BA | 3.047E-01 | 0.481 | 0.519 | 0.115 | 0.115 |
| Vietnam_BA | 6.560E-01 | 0.663 | 0.337 | 0.050 | 0.050 |
| Thailand_IA | 3.194E-01 | 0.635 | 0.365 | 0.065 | 0.065 |
| Qihe_EN | 5.139E-01 | 0.583 | 0.417 | 0.058 | 0.058 |
| Liangdao1_EN | 7.219E-01 | 0.527 | 0.473 | 0.059 | 0.059 |
| Liangdao2_EN | 3.856E-01 | 0.570 | 0.430 | 0.056 | 0.056 |
| Fujian_LN | 1.814E-02 | 0.608 | 0.392 | 0.047 | 0.047 |
| Hanben_IA | 8.145E-01 | 0.688 | 0.312 | 0.031 | 0.031 |
| Gongguan_IA | 5.574E-01 | 0.672 | 0.328 | 0.061 | 0.061 |
| Vanuatu_2900BP_all | 5.854E-01 | 0.638 | 0.362 | 0.061 | 0.061 |
| Daur | 5.628E-01 | 0.883 | 0.117 | 0.032 | 0.032 |
| Xibo | 2.916E-02 | 0.840 | 0.160 | 0.034 | 0.034 |
| Hezhen | 1.081E-01 | 0.905 | 0.095 | 0.033 | 0.033 |
| Nanai | 2.966E-01 | 0.966 | 0.034 | 0.034 | 0.034 |
| Negidal | 1.523E-01 | 1.017 | -0.017 | 0.041 | 0.041 |
| Nivh | 1.268E-01 | 0.979 | 0.021 | 0.036 | 0.036 |
| Oroqen | 9.378E-02 | 0.936 | 0.064 | 0.034 | 0.034 |
| Ulchi | 7.103E-02 | 0.990 | 0.010 | 0.034 | 0.034 |
| Korean | 4.766E-01 | 0.822 | 0.178 | 0.032 | 0.032 |
| Japanese | 1.255E-01 | 0.787 | 0.213 | 0.029 | 0.029 |
| JPT.SG | 9.384E-02 | 0.795 | 0.205 | 0.030 | 0.030 |
| CHB.SG | 1.701E-01 | 0.789 | 0.211 | 0.030 | 0.030 |
| CHS.SG | 3.262E-01 | 0.746 | 0.254 | 0.029 | 0.029 |
| Han_NChina | 3.269E-01 | 0.782 | 0.218 | 0.030 | 0.030 |
| Han_Shanxi | 7.612E-01 | 0.780 | 0.220 | 0.030 | 0.030 |
| Han_Shandong | 3.585E-01 | 0.777 | 0.223 | 0.030 | 0.030 |
| Han_Henan | 3.620E-01 | 0.797 | 0.203 | 0.034 | 0.034 |
| Han_Jiangsu | 6.479E-01 | 0.790 | 0.210 | 0.031 | 0.031 |
| Han_Shanghai | 3.596E-01 | 0.771 | 0.229 | 0.035 | 0.035 |
| Han_Hubei | 3.524E-01 | 0.751 | 0.249 | 0.033 | 0.033 |
| Tujia | 2.355E-01 | 0.761 | 0.239 | 0.030 | 0.030 |
| Han_Chongqing | 3.799E-01 | 0.755 | 0.245 | 0.035 | 0.035 |
| Han_Sichuan | 3.907E-01 | 0.747 | 0.253 | 0.030 | 0.030 |
| Han_Zhejiang | 2.320E-01 | 0.757 | 0.243 | 0.032 | 0.032 |
| Han_Fujian | 3.615E-01 | 0.724 | 0.276 | 0.032 | 0.032 |
| Han_Taiwan | 6.705E-01 | 0.742 | 0.258 | 0.031 | 0.031 |
| Han_Hakka | 4.255E-01 | 0.735 | 0.265 | 0.030 | 0.030 |
| Han_Yunnan | 3.207E-01 | 0.749 | 0.251 | 0.029 | 0.029 |
| Han_Guangdong | 3.999E-01 | 0.712 | 0.288 | 0.031 | 0.031 |
| Han_Guangxi | 4.243E-01 | 0.704 | 0.296 | 0.029 | 0.029 |
| Han_Haikou | 4.678E-01 | 0.721 | 0.279 | 0.028 | 0.028 |
| Monguor | 4.567E-02 | 0.770 | 0.230 | 0.031 | 0.031 |
| Yugur | 1.634E-01 | 0.772 | 0.228 | 0.030 | 0.030 |
| Qiang_Daofu | 1.641E-01 | 0.760 | 0.240 | 0.030 | 0.030 |
| Qiang_Danba | 1.713E-01 | 0.782 | 0.218 | 0.030 | 0.030 |
| Tibetan_Xinlong | 2.645E-01 | 0.759 | 0.241 | 0.031 | 0.031 |
| Tibetan_Gangcha | 1.053E-01 | 0.774 | 0.226 | 0.030 | 0.030 |
| Tibetan_Xunhua | 3.318E-02 | 0.723 | 0.277 | 0.034 | 0.034 |
| Tibetan_Chamdo | 2.783E-01 | 0.770 | 0.230 | 0.030 | 0.030 |
| Tibetan_Lhasa | 3.319E-01 | 0.754 | 0.246 | 0.031 | 0.031 |
| Tibetan_Nagqu | 5.123E-01 | 0.759 | 0.241 | 0.030 | 0.030 |
| Tibetan_Shigatse | 3.908E-01 | 0.724 | 0.276 | 0.031 | 0.031 |
| Tibetan_Shannan | 3.233E-01 | 0.722 | 0.278 | 0.031 | 0.031 |
| Sherpa.DG | 2.050E-02 | 0.713 | 0.287 | 0.039 | 0.039 |
| Naxi | 3.315E-01 | 0.728 | 0.272 | 0.030 | 0.030 |
| Yi | 5.347E-01 | 0.756 | 0.244 | 0.030 | 0.030 |
| Lahu_China | 7.062E-01 | 0.652 | 0.348 | 0.029 | 0.029 |
| Lahu_Vietnam | 7.231E-01 | 0.682 | 0.318 | 0.030 | 0.030 |
| Cong | 6.635E-01 | 0.634 | 0.366 | 0.030 | 0.030 |
| HaNhi | 3.238E-01 | 0.695 | 0.305 | 0.029 | 0.029 |
| Sila | 5.416E-01 | 0.696 | 0.304 | 0.031 | 0.031 |
| She | 3.514E-01 | 0.730 | 0.270 | 0.031 | 0.031 |
| Miao_Hunan | 4.511E-01 | 0.727 | 0.273 | 0.030 | 0.030 |
| Miao_Longlin | 6.374E-01 | 0.687 | 0.313 | 0.033 | 0.033 |
| Miao_Xilin | 3.733E-01 | 0.681 | 0.319 | 0.033 | 0.033 |
| Miao_Rongshui | 1.874E-01 | 0.703 | 0.297 | 0.032 | 0.032 |
| Miao_Sanjiang | 2.764E-01 | 0.681 | 0.319 | 0.033 | 0.033 |
| Miao_Huanjiang | 5.102E-01 | 0.703 | 0.297 | 0.031 | 0.031 |
| Miao_Longsheng | 7.503E-01 | 0.699 | 0.301 | 0.031 | 0.031 |
| Miao_Ziyuan | 1.504E-01 | 0.709 | 0.291 | 0.032 | 0.032 |
| Hmong | 3.666E-01 | 0.692 | 0.308 | 0.031 | 0.031 |
| PaThen | 4.965E-02 | 0.729 | 0.271 | 0.032 | 0.032 |
| Dao | 3.200E-01 | 0.689 | 0.311 | 0.031 | 0.031 |
| Gelao | 2.265E-01 | 0.673 | 0.327 | 0.030 | 0.030 |
| LaChi | 3.818E-01 | 0.666 | 0.334 | 0.033 | 0.033 |
| CoLao | 3.944E-01 | 0.695 | 0.305 | 0.032 | 0.032 |
| Maonan | 3.220E-01 | 0.676 | 0.324 | 0.031 | 0.031 |
| Mulam | 1.359E-01 | 0.691 | 0.309 | 0.031 | 0.031 |
| Dong_Guizhou | 4.464E-01 | 0.704 | 0.296 | 0.031 | 0.031 |
| Dong_Hunan | 3.626E-01 | 0.700 | 0.300 | 0.031 | 0.031 |
| Hlai_Qiongzong | 5.390E-01 | 0.656 | 0.344 | 0.029 | 0.029 |
| Hlai_HO | 2.707E-01 | 0.651 | 0.349 | 0.034 | 0.034 |
| Zhuang_HO | 4.818E-01 | 0.688 | 0.312 | 0.029 | 0.029 |
| Zhuang_Guangxi | 4.656E-01 | 0.689 | 0.311 | 0.028 | 0.028 |
| Zhuang_Yunnan | 2.399E-01 | 0.681 | 0.319 | 0.030 | 0.030 |
| Nung | 4.955E-01 | 0.680 | 0.320 | 0.029 | 0.029 |
| Tay | 3.502E-01 | 0.680 | 0.320 | 0.030 | 0.030 |
| BoY | 9.470E-02 | 0.685 | 0.315 | 0.033 | 0.033 |
| Dai | 2.867E-01 | 0.670 | 0.330 | 0.031 | 0.031 |
| CDX.SG | 4.419E-01 | 0.663 | 0.337 | 0.029 | 0.029 |
| Muong | 6.631E-01 | 0.671 | 0.329 | 0.030 | 0.030 |
| Kinh | 7.350E-01 | 0.663 | 0.337 | 0.030 | 0.030 |
| Vietnamese | 3.438E-01 | 0.678 | 0.322 | 0.030 | 0.030 |
| KHV.SG | 5.111E-01 | 0.664 | 0.336 | 0.028 | 0.028 |
| Dusun | 3.060E-01 | 0.586 | 0.414 | 0.031 | 0.031 |
| Murut | 5.699E-01 | 0.597 | 0.403 | 0.032 | 0.032 |
| Malay | 2.518E-01 | 0.482 | 0.518 | 0.031 | 0.031 |
| Lao | 8.070E-01 | 0.592 | 0.408 | 0.030 | 0.030 |
| Thai | 4.131E-02 | 0.538 | 0.462 | 0.032 | 0.032 |
| Cambodian | 1.697E-01 | 0.515 | 0.485 | 0.031 | 0.031 |
| Mang | 6.467E-01 | 0.615 | 0.385 | 0.030 | 0.030 |
| Htin_Mal | 6.216E-01 | 0.545 | 0.455 | 0.033 | 0.033 |
| Mlabri | 5.319E-01 | 0.498 | 0.502 | 0.039 | 0.039 |
| Ami | 5.141E-01 | 0.642 | 0.358 | 0.032 | 0.032 |
| Atayal | 6.564E-01 | 0.676 | 0.324 | 0.034 | 0.034 |
| Kankanaey | 7.745E-01 | 0.668 | 0.332 | 0.032 | 0.032 |
| Tagalog | 5.997E-01 | 0.592 | 0.408 | 0.031 | 0.031 |
| Malaysia_Jehai.SG | 9.572E-01 | 0.177 | 0.823 | 0.049 | 0.049 |
| Onge.DG | 7.643E-01 | -0.036 | 1.036 | 0.046 | 0.046 |

